# The isolation and characterization of *Taphrina betulina* and other yeasts residing in the *Betula pendula* phylloplane

**DOI:** 10.1101/2021.09.17.460733

**Authors:** Margaretta Christita, Timo P. Sipilä, Kirk Overmyer

## Abstract

The phylloplane is an important microbial habitat and a reservoir of organisms that affect plant health, both positively and negatively. *Taphrina betulina* is the causative agent of birch witches’ broom disease. *Taphrina* are dimorphic, invading theirs hosts in a filamentous form and residing in the host phyllosphere in their non-infectious yeast form. As such, they are widely accepted to be found a resident yeasts on their hosts, even on healthy tissues; however, there is little experimental data to support this. With the aim of exploring the local infection ecology of *T. betulina*, we had isolated yeasts from the phylloplane of birch, using three classes of samples; from infected symptom bearing leaves inside brooms, healthy leaves from branches away from brooms on broom bearing trees, and symptom-free leaves from symptom-free trees. Isolations yielded 224 yeast strains, representing 11 taxa, including *T. betulina*, which was the most common isolate and was found in all sample classes, including asymptomatic leaves. Genotyping with two genetic markers revealed genetic diversity among these *T. betulina* isolates, with seven distinct genotype differentiated by the markers used. Of the 57 *T. betulina* strains, 22 representative strains were selected for further studies and preliminarily characterized, revealing differences in size and the ability to produced compounds with activity to activate the signalling pathway for the plant hormone auxin.

## Introduction

The phyllosphere has been recognized as an important habitat for microorganisms for over a century (Vorholt, 2012). Microbes residing in the phylloplane have various lifestyles and interactions with their hosts ranging from mutualistic symbionts, neutral residents, to pathogens (Vorholt, 2012, Rastogi et al., 2013). Various yeast-like fungi have been reported as resident in different plant compartments including the phylloplane (Fonseca and Inácio, 2006, Begerow et al., 2017, Kemler et al., 2017, Limtong and Nasanit, 2017, Yurkov et al., 2015), some of which can modulate plant health both by directly interacting with the host and by reshaping the microbiome (Agler et al., 2016, Regalado et al., 2020, Brachi et al., 2021).

*Taphrina* are phytopathogenic yeasts causing disease often involving tumour symptoms mostly on woody plant species. These organisms have a dimorphic lifestyle, frequently residing in their host’s phyllosphere for long periods in their haploid budding yeast form and invading their hosts in their pathogenic dikaryotic filamentous form when environmental conditions are favourable (Mix, 1949, Fonseca and Rodrigues, 2011). *Taphrina* species, including *T. betulina* (Kern and Naef-Roth, 1975), are able to produce the plant hormones auxin and cytokinin (Streletskii et al., 2016, Streletskii et al., 2019, Wang et al., 2016, Cissé et al., 2013). Auxin and cytokinin are widely believed to be involved in the many tumour and leaf deformation symptoms caused by various *Taphrina* species, although this is not been tested experimentally. Auxin is also an important adaptation in phyllosphere resident microbes (Fonseca and Inácio, 2006, Kemler et al., 2017) thus may be involved in multiple aspects of lifestyle in *Taphrina* species. The genus *Taphrina* belongs to the order Taphrinales along with its sister genus *Protomyces*, and other genera, all of which are plant pathogens with similar lifestyles and pathogenesis strategies. As members of the ascomycete subphylum Taphrinomycotina, these yeasts possess many ancestral characteristics and thus are of considerable evolutionary interest (Wang et al., 2021, Wang et al., 2019). The genomes of several species in *Taphrinales* are now available opening these organisms to the possibility of molecular studies (Wang et al., 2020, Wang et al., 2019, Cissé et al., 2013, Tsai et al., 2014).

*T. betulina* and several closely related *Taphrina* species (Table 1) are the causative agents of witches’ broom disease on birch (*Betula* spp.), which induce distinctive broom symptoms (Figure 1D) formed from proliferation of axillary buds and shoots around a central infected bud (Mix, 1949, Jump and Woodward, 1994, Fonseca and Rodrigues, 2011, Christita, 2021). Other symptoms include changes in leaves, such as enlargement (but not thickening), chlorosis, leaf curl, leaf spots, premature senescence, and necrosis (Fonseca and Rodrigues, 2011; Mix, 1949). *Taphrina* species, including *T. betulina*, have most often been isolated from hosts with disease symptoms. Exceptionally, some *Taphrina* species, such as *T. inositophila* and *T. kurtzmanii* have been isolated from disease free hosts (Fonseca and Inácio, 2011, Inácio et al., 2004).

**Table 1:** *Taphrina* species infecting *Betula* species^a^.

| Species | Spore sizes (µm) | Species observed on |
| --- | --- | --- |
| *Taphrina betulicola* | spores 3.5 x 3 | *Betula emani* |
| *T. betulina*<sup>b</sup> | Ascospores 4.5-6.5 x 4-5.5<br>Blastospores 3.5-6 x 2-4.5 | *B. aurata*, *B. carpatica*, *B. intermedia*<sup>c</sup>, *B. nana*, *B. pendula*, and *B. pubescens* |
| *T. nana*<sup>d</sup> | Ascospores 3.5-6 x 3.5-5 | *B. emani*, *B. intermedia*, *B. japonica*, *B. nana*, *B. pendula*, and *B. pubescens* |
| *T. americana* | Spores 4-6 x 3.5-5 | *B. fontinalis*, *B. lutea*, and *B. papyrifera* |
| *T. boycei* | Ascospores 4-5 x 3.5-4 | *B. fontinalis* and *B. occidentalis* |
| *T. carnea*<sup>e</sup> | Ascospores 5-6 x 3-4<br>Blastospores 3-6 x 2-4 | *B. fruticose*, *B. glandulosa*, *B. humilis*, *B. intermedia*, *B. lutea*, *B. nana*, *B. papyrifera*, *B. pendula*, and *B. pubescens* |
| *T. bacteriosperma* | Blastospores 3-6 x 1-2 | *B. glandulosa*, *B. intermedia*, *B. nana*, and *B. pubescens* |
| *T. betulae* | Ascospores 4-6 x 3.5-5 | *B. intermedia*, *B. medwediewi*, *B. pendula* var. *purpurea*, *B. pubescens*, and *B. turkestanica* |
| *T. flava* | Blastospores 5-6 x 5-5.5 | *B. papyrifera* and *B. populifolia* |

<sup>a</sup>Compiled from Mix (1949) and Fonseca and Rodrigues (2011). <sup>b</sup>Includes the species *Exoascus betulinus*, *E. turgidus*, *T. turgida*, *T. willeana*, *T. lapponica*, *E. lapponicus*, *T.* *lagerheimii*, and *T. splendens* shown by Mix (1949) to be synonymous with *T. betulina*. <sup>c</sup>*B.* *intermedia* is a hybrid (*B. nana* x *B. pubescens*) <sup>d</sup>*T. nana* has an ITS sequence and PCR fingerprint identical to *T. betulina* (Rodrigues and Fonseca, 2003) and includes the synonymous species *T. alpina* (Mix 1949). <sup>e</sup>*T. carnea* has an ITS sequence and PCR fingerprint identical to *T. betulina* (Rodrigues and Fonseca, 2003) and includes the synonymous species *T. janus* and *T. lata* (Mix 1949).

**Figure 1.**
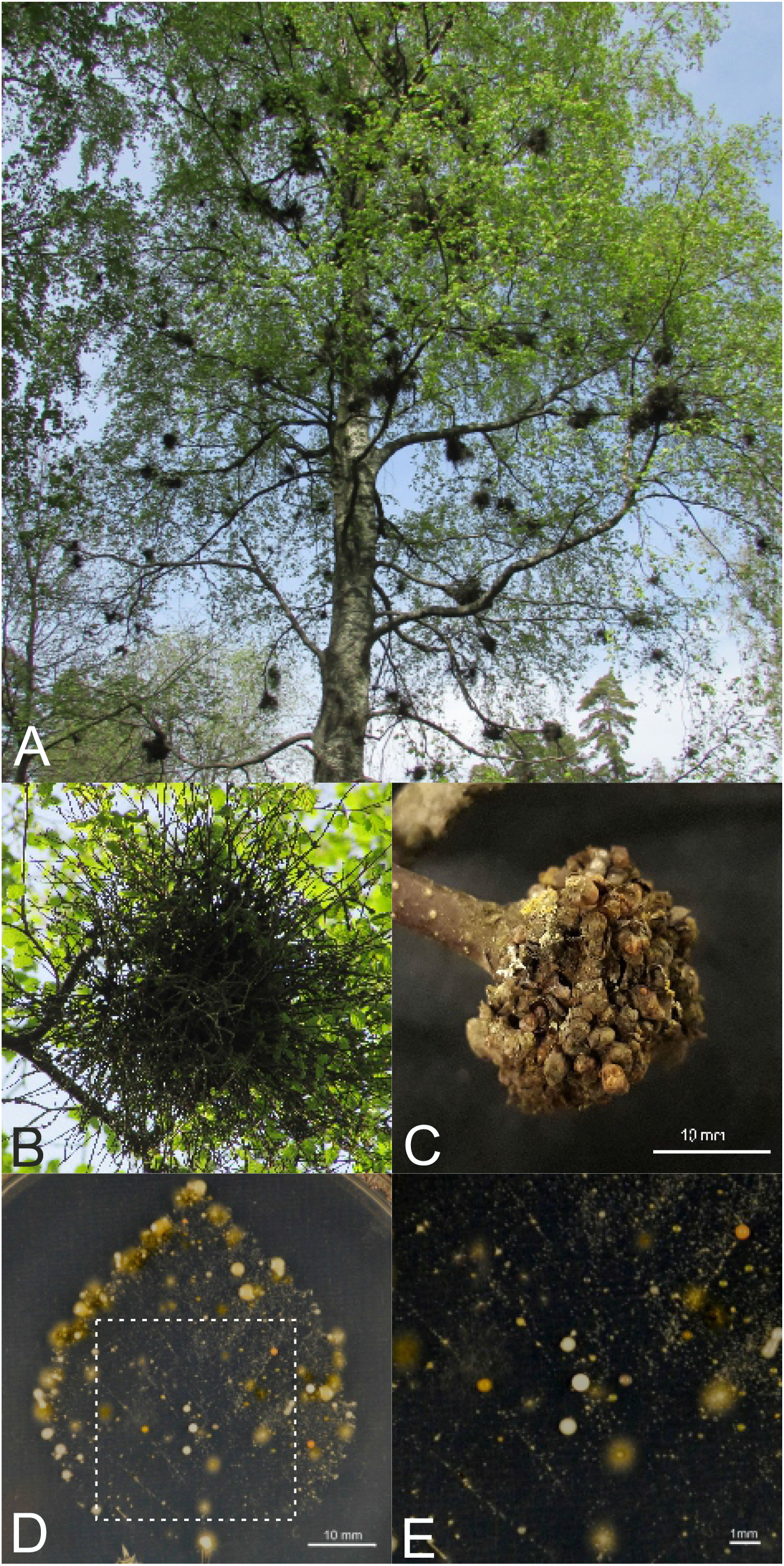
Witches’ broom disease symptoms and leaf press culture. (**A**). Typical witches’ broom symptoms on a heavily infected birch tree. **(B**). Detail of a typical broom with elongated shoots. **(C)**. Detail of an atypical broom symptom in which the central woody tumour is covered in buds that have not elongated into shoots. Size bar = 1cm (**D**). Birch leaf press culture demonstrating the presence of yeasts in the phylloplane of *B. pendula*, which have been cultivated for 14 days on 0.2 x PDA medium. Size bar = 1cm (**E**) Close up details of the area marked with a box in (D) showing colonies with a typical yeast morphology, some of which are consistent with known colony morphology and cream colour of *T. betulina*. Size bar = 1mm

The impact of witches’ broom disease is generally underappreciated and often regarded a curiosity more than a damaging disease (Price and Macdonald, 2012). However, the few studies that have addressed this issue illustrate the need to re-evaluate this view. A negative impact of witches’ broom disease on *B. pubescens* has been demonstrated, showing decreased stem quality, vigour, and growth (Spanos and Woodward, 1994). In *T. betulina* infected *B. maximowicziana* leaves, reduced photosynthesis and dark respiration, chlorosis, and leaf loss were reported (Koike and Tanaka, 1986). The presence of witches’ brooms in birch trees was shown to cause rapid death of the branches where they form and to negatively alter crown architecture (Kostina et al., 2015). Consequently, this disease may result in production losses under short rotation forestry programs (McKay, 2011).

The genomes of several birch species are now available (Salojarvi et al., 2017, Wang et al., 2013). The utility of birch as a model woody forest species has been demonstrated, including use of molecular studies and induced rapid flowering for forward genetic studies (Alonso-Serra et al., 2020, Alonso-Serra et al., 2019). These developments make birch an attractive model for the study of microbial interactions with long lived forests species.

Several yeast genera have been reported as common members of phylloplane communities on multiple plant species, such as; *Rhodotorula, Cryptococcus, Sporobolomyces*, and *Diozegia* (Fonseca and Inácio 2006). Previous studies have addressed the fungal endophytes (Helander et al., 2007) and fungal phylloplane residents of birch (Helander and Rantio-Lehtimaki, 1990, Helander et al., 1993, Nguyen et al., 2017), Only one of these studies (Nguyen et al., 2017) has identified yeasts in the Taphrinales (OTU2 and OTU24) on birch, but these were not identified to the species level. Thus, the basic ecology of *T. betulina* on birch remains uncharacterized. Here we isolate and identify resident yeasts from the *B. pendula* phylloplane, with the goal of investigating the local ecology of *T. betulina* strains.

## Materials and methods

### Sampling, isolation, and cultivation of yeasts

Leaves of birch (*Betula pendula* Roth) were sampled from several locations around eastern Helsinki, Finland (Table 2), placed into sterile 50 ml centrifuge tubes (Sarstedt; www.sarstedt.com) and stored at +4C prior to further processing. Leaf press cultures were done by placing birch leaves on the surface of a plate containing 0.2x potato dextrose broth (PDB; BD Difco; www.fishersci.com) with 15g/l agar (PDA), covered with sterile filter paper, and pressed with gloved fingertips to transfer phylloplane microbes to the media. Leaf press culture plates were cultivated one to two days at 21°C and then stored at 4°C for two weeks to promote colony colour formation. For yeast isolation, three samples were collected for type I samples, which were symptomatic leaves (harvested from inside brooms) from symptomatic trees, five samples for type II samples, which were asymptomatic leaves from symptomatic trees, and two samples were collected for type I samples, which were asymptomatic leaves from asymptomatic trees. Leaf samples were gently washed three times in sterile ultrapure water. Leaves were placed in sterile 50ml centrifuge tubes with 5 ml of sterile ultrapure water with 0.025% Silwet-L77, then spores were released by vigorous agitation with a vortex mixer. A dilution series (10^−1^ to10^−3^) of the leaf wash solution was plated on 0.2x PDA plates and cultivated for one week at 21°C under a 12 hr light cycle in a controlled environment chamber (Model MLR-350, Sanyo, uk.vwr.com). Colonies were picked after three, five, and seven days, and isolated by two rounds of streaking onto fresh 0.5xPDA plates supplemented with 100 ug/ml Ampicillin. An equal number of plates made for each sample and the number of colonies picked represented the number and diversity of colonies present on the plates. Wash solutions and isolated strains were placed in 50% sterile glycerol at −80°C for long term storage. For short term storage cells from each strain were inoculated into 200 µl sterile water in a 96 well plates to facilitate high-throughput inoculation, cultivation, and DNA isolation.

**Table 2.**
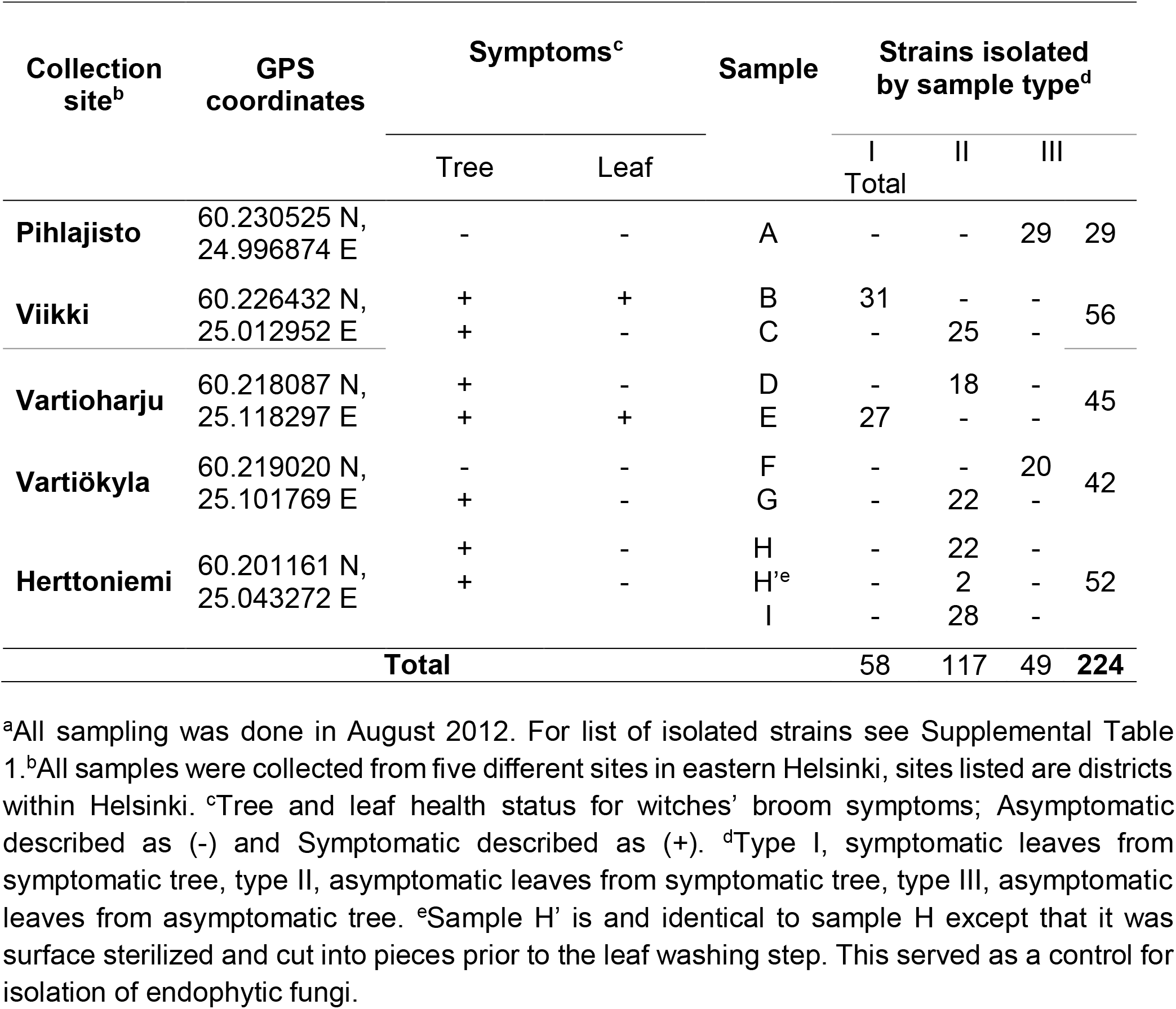
Sampling sites and leaf samples collected^a^.

### *DNA extraction* and yeast identification

Following cultivation in 2 ml 0.2 × PDB for 5 days, cells were harvested by centrifugation (5 min x12,000 g) for DNA isolation as in (Looke et al., 2011). Briefly, cells were resuspended in one volume lysis buffer (100 mM Tris-HCl pH8, 50 mM EDTA, 500 mM NaCl), then vortexed for 3 min with 0.3 g glass beads and 200 μl phenol/chloroform/isoamyl alcohol. After adding 200 μl TE, samples were briefly vortexed, then centrifuged for 5 min. The aqueous layer was transferred to a clean 2 ml tube, ethanol precipitated, pelleted, resuspended in 0.4 ml TE buffer, treated with DNase-free RNase A (5 min at 37 °C), ethanol precipitated, and dissolved in 100 μl TE. Yeast strains were identified based on sequences of their internal transcribed spacer (ITS) region of the nuclear rRNA locus. ITS PCR products were amplified from yeast genomic DNA with the ITS3 (5-CTTGGTCATTTAGAGGAAGTAA-3) and ITS4 (5-TCCTCCGCTTATTGATATGC-3) primers as described in (Wang et al., 2016). Morphologically similar yeasts with identical ITS PCR product lengths underwent cleaved amplified polymorphic sequence (CAPS) analysis by digesting the ITS PCR product with TaqI (Thermo scientific) according to the manufacturer’s instructions and separating the fragments on a 3% agarose gel. This ITC (ITS Taq 1 CAPS) marker identified 17 variants ITC-A to ITC-U. Isolates were binned based on identical ITS restriction patterns and a representative of each group was sequenced using ITS1 (5-GGAAGTAAAAGTCGTAACAAGG-3) and ITS4 primers. Prior to sequencing, primers remaining from PCR amplification were removed by treatment with ExoSAP (Exonuclease I, Shrimp Alkaline Phosphatase; Thermo Scientific; https://www.fishersci.fi/). Assembled complete ITS regions (ITS1 - 5S rRNA - ITS2) were queried against sequences of known fungi using the Basic Local Alignment Search Tool (BLAST) at the NCBI (https://www.ncbi.nlm.nih.gov/).

### Morphological characterization

*Taphrina* strains were grown at 0.2x PDA media using streak technique on 9 cm diameter petri dish and were cultivated at constant 23°C in a controlled environment chamber (Sanyo MLR-350; www.sanyo-biomedical.co.uk) for 3 days. Microscopic cell investigation including length, width, and shape were performed on three-day-old cultures. Cell images were captured by LEICA 2500 microscope with camera LEICA DFC490. Cell length and width were measured with imageJ software (imagej.nih.gov/ij/).

### Indolic compound production and auxin response assay

Salkowski reagent colorimetric assay (Glickmann and Dessaux, 1995) was used to detect indolic compounds including auxin. Two different liquid media, YPD, (yeast extract, peptone, dextrose) and YPD +0.1% tryptophan, were used to cultivate 22 *T. betulina* strains for five days. Yeast cells were sedimented by centrifugation (12000 rpm, 20°C for 5 min) and 0.5 ml culture supernatant mixed with 0.5 ml Salkowski reagent, then A_530_ was measured. The experiment was performed in three independent biological replicates, each with three technical replicates.

For visualizing the activation of the plant auxin transcriptional response, *Arabidopsis thaliana (*hereafter referred to as *Arabidopsis)* Col-0 accession plants transgenically bearing the auxin-responsive promoter::reporter system, DR5::GUS (β-glucuronidase) were used. Seeds were sown on 0.5x MS agar medium in 6 well plates, 5 plants per well, then stratified in the dark at 4°C for two days, then transferred to a growth chamber with 12/12hr light/dark cycle at 24°C. Ten-day-old DR5::GUS seedlings were treated overnight with 1 ml filtered culture supernatants from five-day-old *T. betulina* cultures (YPD and YPD with 0.1% L-tryptophan). Positive controls were treated with 5 μM IAA, negative control with filtrate from uncultivated medium. GUS staining solutions were prepared with 1 mM 5-bromo-4-chloro-3-indolyl b-D-glucuronide dissolved in methanol, 5 mM potassium ferricyanide and 5 mM potassium ferrocyanide in 50 mM sodium phosphate buffer (pH 7.2). For histochemical staining, seedlings were fixed with ice-cold 90% acetone for 1 h, washed two times with ice-cold wash solution (36 mM Na_2_HPO_4_, 14 mM NaH_2_PO_4_, pH 7.2), 30 min for each wash. Seedlings were vacuum infiltrated for 5 min and kept at room temperature in GUS staining solution. Stained seedlings were washed two times with absolute ethanol and stored in 70% ethanol.

## Results

### Sample Collection and Yeast Isolation from Birch

*Taphrina species*, including *T. betulina*, have been most commonly isolated from symptomatic host leaves (Mix, 1949) and have only infrequently been isolated from healthy tissues (Fonseca and Inácio, 2011, Inacio et al., 2004). To assess if *Taphrina*-like yeasts could be isolated from the phyllosphere of asymptomatic birch leaves, leaf press cultures were performed using birch leaves collected in the field from asymptomatic trees. Following cultivation using media and conditions that favoured yeasts, a high diversity of yeast colonies were apparent (Figure 1D). The leaf press culture revealed a large diversity of yeast in the birch phylloplane, including yeasts coloured with white, beige, and orange to pink pigments, some of which are consistent with *T. betulina* (Figure 1 D-E).

We isolated birch associated yeasts with the aim of obtaining new strains and probing the ecology of *Taphrina betulina* and other yeasts within and between infected and healthy individual host trees. A total of nine leaf samples (A-I) were collected from both symptomatic and asymptomatic trees at five independent sites in eastern Helsinki (Table 2). Samples were classified into three types based on the tree’s health characteristics; type I is symptomatic leaves from a symptomatic tree, type II is asymptomatic leaves from a symptomatic tree, and type III is asymptomatic leaves from an asymptomatic tree. Witches’ broom disease on birch can manifest in distinctive types of broom symptoms; typical brooms have multiple elongated shoots that have grown from many ectopic axillary buds, which form around the primary infected bud, to form a central tumour (Figure 1B). Occasionally, infected branches are found with a central tumour covered in buds that have not elongated into shoots (Figure 1C). There is also a continuum of variation in the length of the shoots, in between these two extremes. To address the possibility that different yeasts may be associated with these various structures, the phenotype of collected type I samples was classified as elongated brooms (EB), short elongated brooms (SE), or tumour-like (TL), while samples originating from host types II and III, which lacked broom symptoms, are classified as no broom (NO) (Tables S1 and S2).

A total of 224 yeast strains were isolated from the birch phyllosphere (Table 2); with 58, 177, and 49 strains coming from type I, II, and III samples, respectively. Of the 224 strains obtained (Table S1), 116 strains were identified in 11 taxa (Figure 2; Table 3); seven defined at the species level, namely *Taphrina betulina, Rhodotorula phylloplana, Rhodotorula bacarum, Rhodotorula laryngis, Cryptococcus wieringae, Cryptococcus tephrensis*, and *Nakazawaea hosltii*; and four defined only to higher level classifications, specifically, *Kuraishia* sp., *Udinomyces* sp., order *Myriangales*, and class *Tremellomycetes* (Figure S2A; Table 3). The remaining 51 strains could not be unambiguously identified. The number of isolates was similar in all samples (Table 2), with two exceptions. Sample I’ was not a true independent sample, but a control that was identical to sample I, except that it was surface sterilized and sliced prior to the leaf wash step. This was done to control for endophytic yeasts. The low number of isolates in this sample suggests that the overwhelming majority of yeasts isolated in this study when epiphytic (phylloplane residents). The number of leaf samples for each sample type varies (3, 5, and 2 for types I-III, respectively; Table 2). In order to compensate for this sampling bias and facilitate comparisons between samples of differing levels of witches’ broom disease symptoms, the normalized number of isolates (average number of isolates per sample type) are presented (in parentheses; Table 3).

**Table 3.** Yeast isolated from birch phylloplane^a^.

| Species | Type I<sup>c</sup> | Type II<sup>d</sup> | Type III<sup>e</sup> | Total |
| --- | --- | --- | --- | --- |
| ***T. betulina* classifications<sup>b</sup>** |  |  |  |  |
| ITC-C RGR-0 | 1 (0.5) | 11 (2.2) | - |  |
| ITC-C RGR-1 | - | - | 1 (0.5) |  |
| ITC-C RGR-2 | - | 1 (0.2) | - |  |
| ITC-D RGR-0 | 8 (4) | 1 (0.2) | 1 (0.5) |  |
| ITC-D RGR-1 | - | 1 (0.2) | - |  |
| ITC-D RGR-2 | 14 (7) | 11 (2.2) | 4 (2) |  |
| ITC-D RGR-3 | - | 3 (0.6) | - |  |
| ***T. betulina* total** | **23 (11.5)** | **28 (5.6)** | **6 (3)** | **57** |
| **Other Species** |  |  |  |  |
| ***Novel Myriangiales* sp.** | - | 5 (1) | 19 (9.5) |  |
| ***Rhodotorula phylloplana*** | 4 (2) | 28 (5.6) | 4 (2) |  |
| ***Rhodotorula bacarum*** | -- | 29 (5.8) | 13 (6.5) |  |
| ***Rhodotorula laryngis*** | 1 (0.5) | - | - |  |
| ***Cryptococcus wieringae*** | - | - | 2 (1) |  |
| ***Cryptococcus tephrensis*** | 4 (2) | - | - |  |
| ***Kuraishia* sp.** | 1 (0.5) | - | - |  |
| ***Nakazawaea holstii*** | 1 (0.5) | - | - |  |
| ***Udeniomyces* sp** | 1 (0.5) | - | - |  |
| ***Tremellomycetes*** | - | 4 (0.8) | - |  |
| **Unidentified** | **23 (11.5)** | **23 (4.6)** | **5 (2.5)** |  |
| ***Other species total*** | **35 (17.5)** | **89 (17.8)** | **43 (21.5)** | **167** |
| **Total species** |  |  |  | **224** |

**Figure 2.**
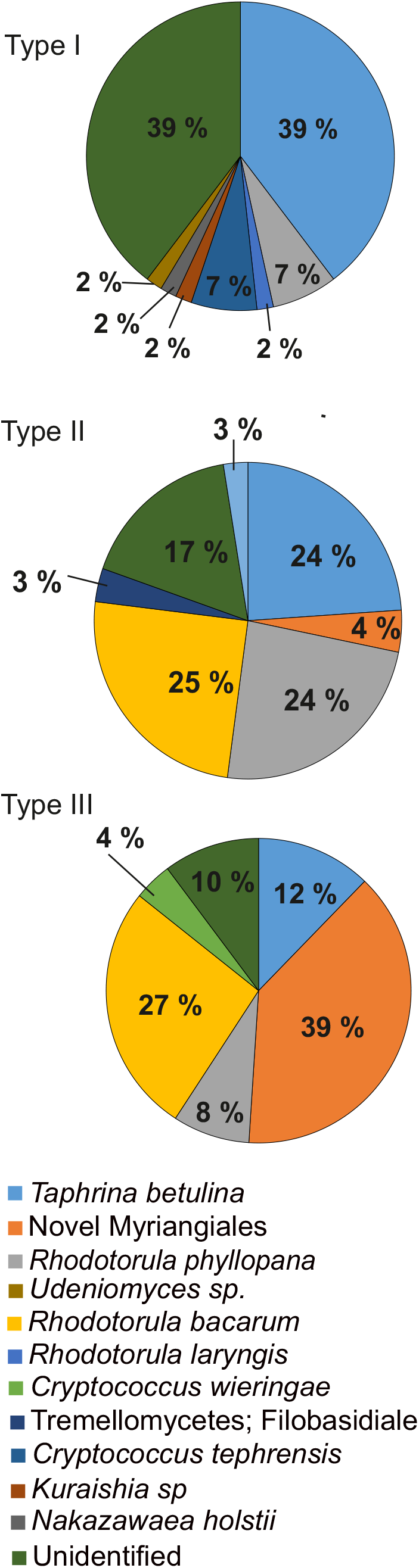
Identification of yeasts isolated from the phylloplane of birch leaves. The identified yeasts isolated from the birch phyllosphere from three sample types (I, symptomatic leaf from a symptomatic tree; II, asymptomatic leaf from a symptomatic tree; III), are presented as percent of the total isolates in each of the three respective samples.

The dominate isolate, with a total of 57 strains (26% of all isolates) was *T. betulina* (Table 3). *T. betulina* was present on all sample types, indicating the species can be isolated from disease-free birch leaves. The further analysis of *T. betulina* strains is discussed below. Other yeast strains found in large numbers were *Myriangiales, R. phylloplana* and *R. bacarum*. Notably, although the level of diversity of yeast present was fairly constant, some differences in the yeast species isolated were observed between the samples differing in their levels of witches’ broom disease symptoms. *Myriangiales* was dominant on type III host samples. *R. phylloplana* was found on all three host types, while *R. bacarum* was prevalent on type II and III host samples. In spite of its presence in high numbers on healthy leaves, *R. bacarum* is absent from leaves in witches’ brooms, suggesting that *T. betulina* alters the phyllosphere microbiome within diseased tissues. A general trend was seen where the total number of isolates for species other than *Taphrina* decreases in increasingly diseased tissue, while the number of *T. betulina* isolates shows the opposite trend (Fig. 2; Table 3). Taken together these results are consistent with the concept of dysbiosis, an imbalance or alteration of microbial communities in diseased tissues.

### *T. betulina* Identification and Characterization

In the order Taphrinales, which includes the genera *Taphrina* and *Protomyces*, sequences of the nuclear rRNA internal transcribed sequences (ITS) are known to give phylogenetic resolution only to the genus level, or to the species level for some taxa (Rodrigues and Fonseca, 2003, Wang et al., 2021, Wang et al., 2019). ITS sequences are sufficient to identify *T. betulina*, however, there are currently no lineage specific secondary phylogenetic markers able to resolve different strains of this species. Accordingly, ITS CAPS analysis of the *T. betulina* strains isolated here only identified two different banding patterns (ITC-C and ITC-D; Figure S3; Table S1). The genotype ITC-D has an ITS sequence identical to *T. betulina*. Also, *T. betulina, T. nana*, and *T. carnea* have identical ITS sequences (Supplemental Figure S3F) (Rodrigues and Fonseca, 2003). These *Taphrina* were isolated as distinct birch species, but later shown by molecular analysis to be conspecific (Rodrigues and Fonseca, 2003). We utilized these and other closely related *Taphrina* species to develop a marker that can differentiate between strains of *T. betulina*. Using the assembly of *T. deformans* genome (Cissé et al., 2013) and our unpublished *T* .*betulina* genome data (PRJNA188318), two pairs of adjacent conserved housekeeping genes (*Rco1*-*Gyp7* and *Sad1-Rax1*) were identified to have conserved synteny between *T. betulina* and *T. deformans*. Two sets of outward facing primers were designed within conserved domains of the gene pairs, allowing amplification of the polymorphic intergenic regions (Supplemental Figure S3B). Both primer pairs gave PCR products of the expected size with *T. betulina* genomic DNA; however, amplification with *T. deformans* DNA only occurred with the primers spanning the *Rco1* and *Gyp7* genes, suggesting they were more highly conserved and suited for marker development (Supplemental Figure S3C). Further tests demonstrated that these primers were able to amplify genomic DNA from all the tested birch associated *Taphrina* species, except *T. americana* (Supplemental Fig 3D). This PCR product digested with RsaI was used as an RFLP marker, which was able to distinguish between *T. deformans*, the related alder pathogen *T. robinsoniana*, and the three *T. betulina* strains with identical ITS sequences (Supplemental Fig. 3D). This marker was named RGR1 (*Rco1 Gyp7* RsaI) and was used to characterize *T. betulina* isolates. The ITS Taq1 CAPS (ITC) marker identified two variants (ITC-C and ITC-D) within the isolated *T. betulina* strains (Table 3; Supplemental Table 1). These were further genotyped using the RGR1 marker, which identified four variants (RGR-0, no PCR product; and RGR-1 to RGR-3, banding pattern variants I-III, respectively). Taking both the ITC and RGR markers into account, seven distinct genotypes were identified within our *T. betulina* isolates (Table 3; Supplemental Table 1).

In table 3, a stable level of strain diversity across all host sample types was seen for *T. betulina*. For the other isolated yeast species diversity was higher in the leaves sampled in brooms, as compared to healthy leaves from both healthy or symptom bearing trees (Table 3). In all cases the strains/species present were different.

The *T. betulina* strain ITS-D RGR-2 was dominant in all host types and followed the general trend of increasing levels in more diseased tissues. There were no clear differences in the *T. betulina* strains present on samples with different broom morphologies (Supplemental Table 2). The distribution of strains was quite varied with some specific to only diseased trees and others present on all trees. This suggests that the length of shoot elongation within brooms caused by *T. betulina* is more likely dependent on the host genotype, or other unknown factors, rather than the *T. betulina* strain present.

We selected 22 of 57 strains of *T. betulina*, which were representative of the diversity in the collection in terms of their genotype and sample origin (Table 4). Half of the 22 selected strains originated from type II, while type I was 32% and type III was 18% (supplemental figure 2) and all genotype were represented (Table 4). This strain collection was used for further identification and characterization. First the morphology of the selected strains was characterized. Colonies of 22 selected strains streaked on 0.2x PDA plates exhibited variation in colour from pale pink to peach (Supplemental Fig. 4, inset) while yeast cells did not exhibit any notable morphological differences (Supplemental Fig. 4). The average yeast cell sizes of all strains taken together were within the range of 3.17-3.81 × 4.75-5.79 µm (Supplemental Table S3), which were generally in agreement with the previously published results for *T. betulina*; 2.4-4 × 2-4 µm (Mix, 1949), 4.5-6 × 4-5 µm (Bacigalova, 1997), 4.5-5x 6.7-8.2 µm (Fonseca and Rodrigues, 2011).Size variation between individual strains within the collection was observed in plots of the average length and width (Figure 3; Supplemental Table S3). Statistical analysis using ANOVA with Tukey’s posthoc test showed that strains 11, 20, 58, and 129 had significant differences to other strains (Supplemental Table S3; Figure 3). There was one common feature to all these strains -- they originated from healthy leaves (Table 4); strains 11, 20, and 129 were from completely healthy trees (type III samples), and strain 58 from healthy leaves on a broom-bearing tree (type II sample). All *T. betulina* strains genotyped as ITC-D RGR-2 grouped together in the size plot indicating they have similar cell sizes.

**Table 4.**
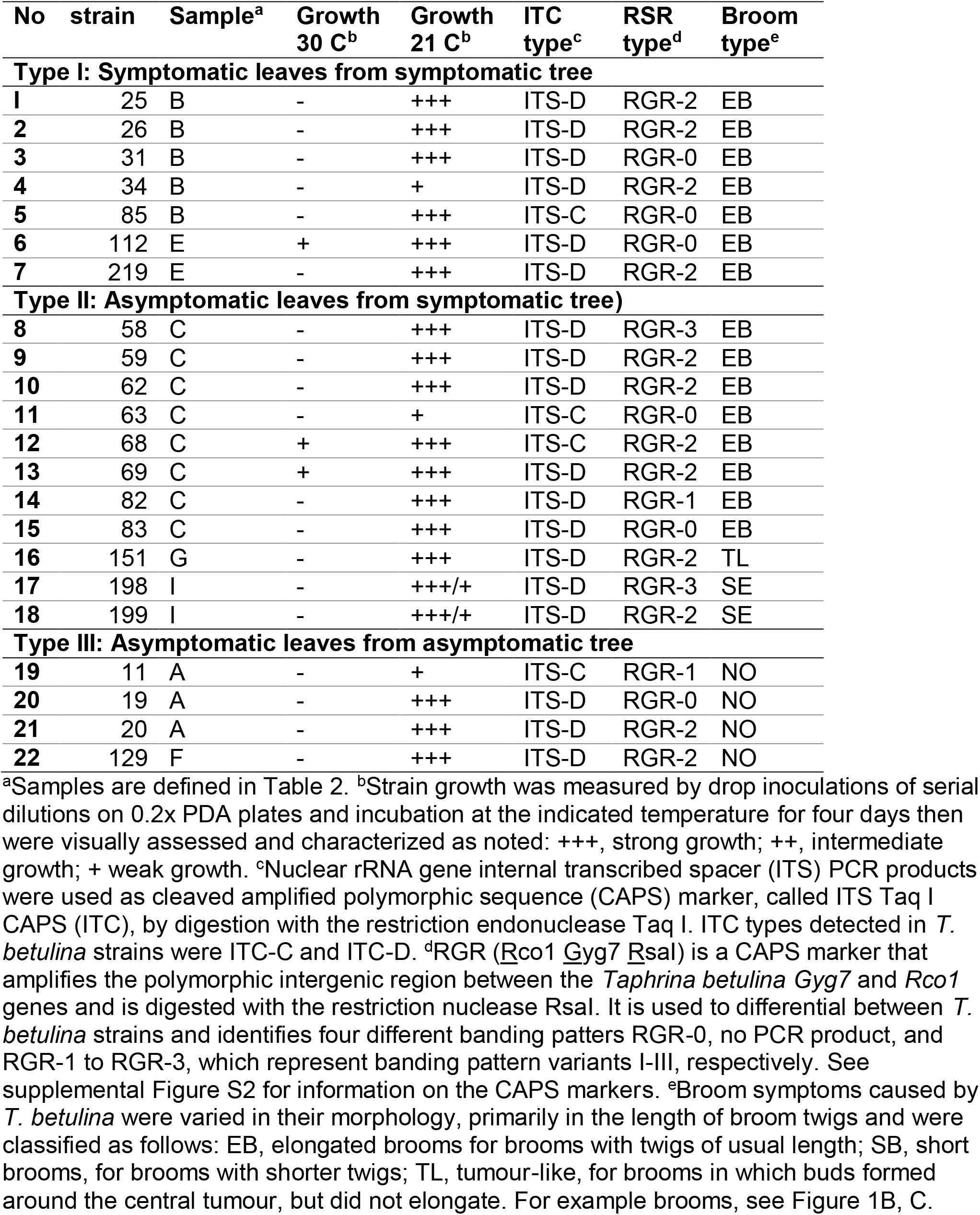
Representative *Taphrina betulina* strains selected for in depth analysis.

**Figure 3.**
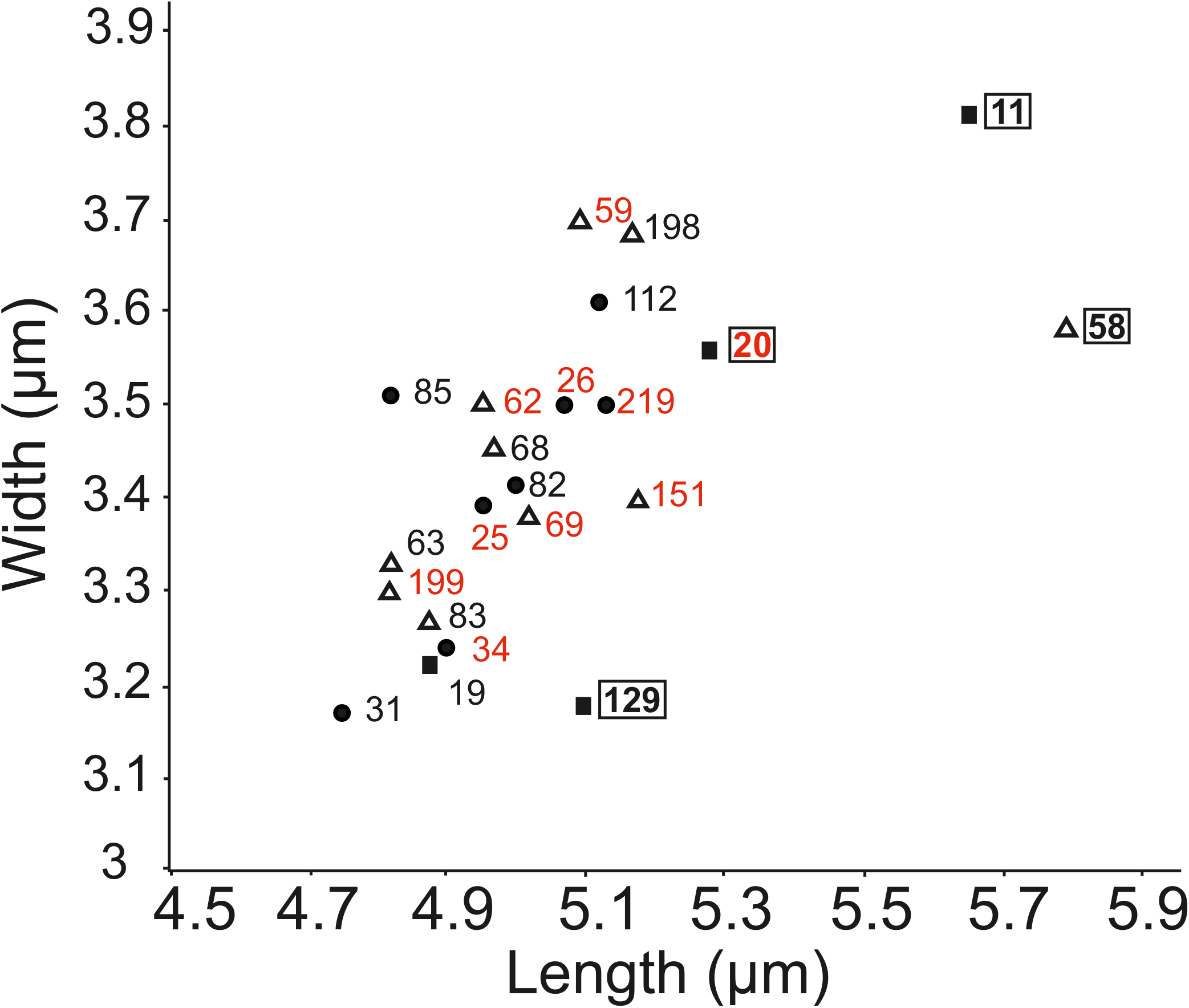
Scatter plot of *T. betulina* strain cell size. Cell sizes for the 22 selected *Taphrina betulina* strains. Each strain number is given next to the data points. Sample type from which the strain originated is indicated by the data point shape as follows; circles, type I samples (symptomatic leaves from symptomatic trees); triangles, type II samples (asymptomatic leaves from symptomatic trees); squares, type II samples (asymptomatic leaves from asymptomatic trees). Numbers enclosed in a box indicate strains whose sizes exhibited statistically significant differences (see Supplemental Table 3) and number depicted in red represent strains of the genotype ITC-D RGR-2.

Production of the indolic plant hormone auxin is a common feature of *T. betulina* (Kern and Naef-Roth, 1975) and is known to have roles in phylloplane residency, pathogenesis, and possibly in tumour symptom formation (Fonseca and Inácio, 2006, Fu and Wang, 2011, Spaepen and Vanderleyden, 2011, Kemler et al., 2017). The Salkowski assay was utilized to quantify production of indolic compounds, used here as a proxy for auxin production. The selected *T. betulina* strains were cultivated in two different media; YPD and YPD+0.1% tryptophan, which is the biosynthetic precursor of indole acetic acid (IAA), the most important plant auxin species. All strains were able to produce indolic compounds in both media (Fig. 4). Indolic compound production was higher for all strains when cultivated in YPD supplemented with tryptophan, as compared to YPD medium alone (Fig. 4). There were marked differences in indolic compound production between strains (Fig. 4a); however, this did not correlate with the sample type of origin, with both high and low producers found in individuals from all three sample types.

**Figure 4.**
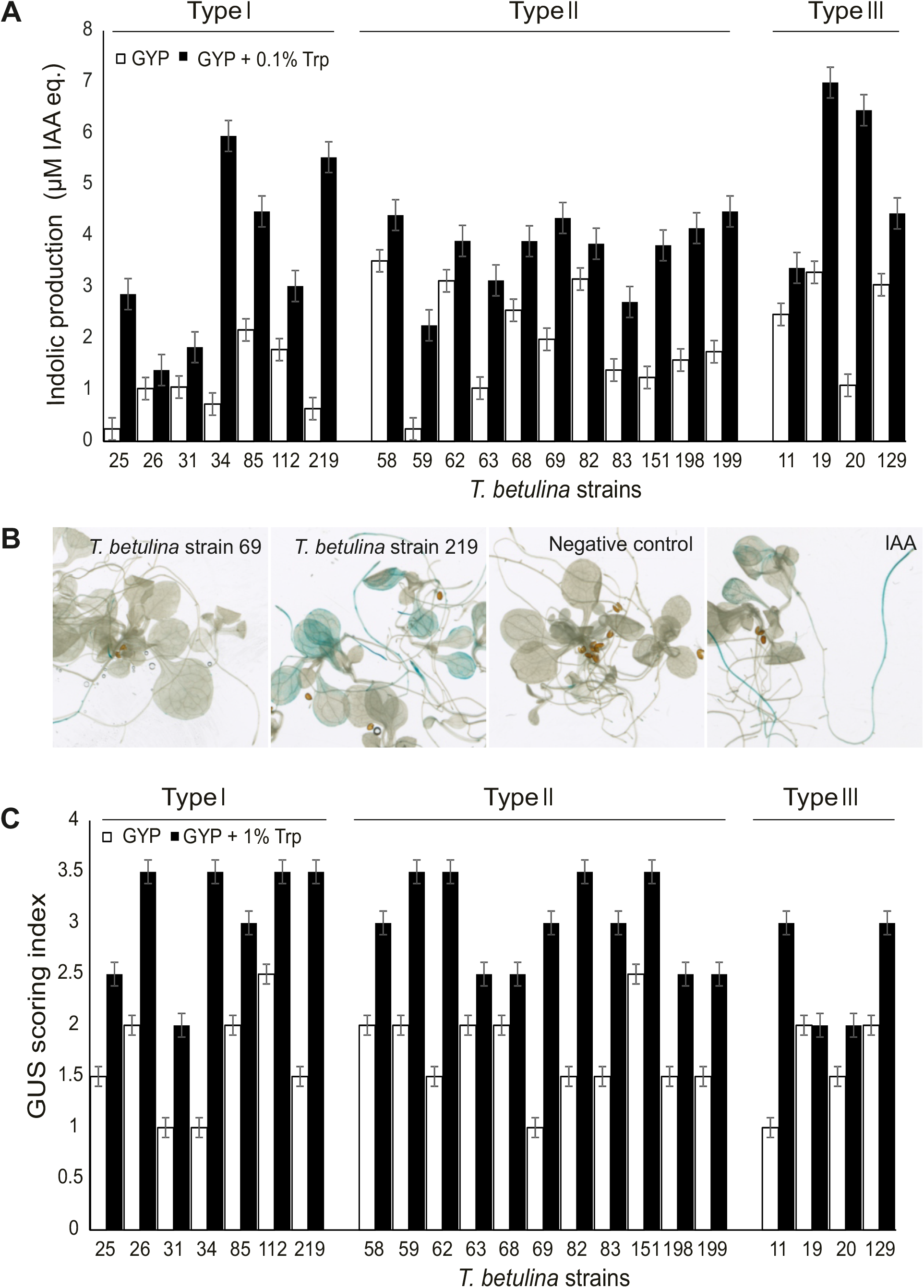
Activation of plant auxin response by *Taphrina betulina* culture filtrates. **(A)** Indolic compound production was used as a proxy for auxin production and was assayed spectophotometrically using the Salkowski assay. *T. betulina* strains (22 total) were cultivated in YPD (yeast-extract, peptone, dextrose) media and with and without 0.1% tryptophan. Three independent biological repeats each with three technical replicates were done for each individual strain. Production levels were calibrated according to a standard curve with the auxin, indole acetic acid (IAA) and are expressed as IAA equivalents. **(B)** The activation of *Arabidopsis* auxin transcriptional response by *T. betulina* culture filtrates was monitored as expression of the auxin responsive DR5 promoter fused to the β-glucuronidase (GUS) reporter gene. Two-week-old *in vitro* grown DR5::GUS *Arabidopsis* seedlings in 0.5 MS media were treated with filtered supernatants of five-day-old yeast culture in YPD and YPD+0.1% tryptophan for 24 hours. Representative results are shown here: light root tip staining was seen in strain 69 cultured in YPD, YPD+ 0.1% tryptophan was used as a negative control. A typical strong response was seen in strain 219 cultured in YPD +0.1% tryptophan and treatment with 5μM IAA was used as a positive control. For full results from all strain see Supplemental Figure S5. **(C)** DR5::GUS staining results in Supplemental Figure S5 were visually scored using an arbitrary GUS scoring index on a scale from 0 to 4. The representative responses shown in (B) were used for reference; where 0 is defined by the negative control and 4 by strain 219. Results from two independent biological repeats were scored and averaged.

In order to gain further evidence for possible differences in auxin production by *T. betulina* strains, a plant based auxin-response reporter system was used. *Arabidopsis* seedlings transgenically bearing the auxin responsive DR5 promoter fused to the β-glucuronidase (GUS) reporter (DR5::GUS) were treated with yeast culture supernatants, then GUS histochemical staining was used to visualize tissues exhibiting activation of the plant auxin transcriptional response. Representative strains exhibiting low and high levels of GUS staining (strains 69 and 219, respectively) are shown (Figure 4b) along with the negative control (uncultivated media) and positive control (5 µM IAA). Photos of all treatments (Supplemental Figure S5) were visually scored on an arbitrary scale from 0 (no staining) to 4 (high staining) and are summarized in Figure 4c.Treatment with culture supernatants of *T. betulina* strains resulted in activation of plant auxin response, to varied levels (Figure 4c; Supplemental Figure S5). Strains isolated from type III samples exhibited lower staining levels suggesting a lower capacity to produce auxin. However, this was only apparent with culture filtrates from cells cultivated with exogenous tryptophan; there were no remarkable differences among samples cultivated in YPD, which represents more natural conditions (Figure 4c; Supplemental Fig. 5).

## Discussion

*Taphrina* are enigmatic and little studied phytopathogenic yeast-like fungi. Many of the basic details of *Taphrina* biology remain unknown or are little studied with modern methods. There is a large body of literature from previous studies of *Taphrina* species (Fonseca and Rodrigues, 2011, Mix, 1949). However, most of this work is quite outdated and even the known aspects of *Taphrina* biology warrant revalidation with modern methods. Here we have addressed the local distribution of *T. betulina* strains with emphasis on the distribution between and within birch branches presenting differing levels of symptoms. Such work had not previously been undertaken and has implications for several aspects of *T. betulina* biology.

### Isolation of *T. betulina* from uninfected tissues

*Taphrina betulina* is typically isolated by the spore drop method from infected leaves with ascogenous cells breaking though the surface (Mix, 1949, Tavares, 2004). In this study, the use of modern high-throughput isolation utilizing rapid PCR based molecular identification methods has facilitated the isolation of *T. betulina* from both symptom bearing material and healthy leaves. The molecular markers utilized in this work allowed differentiation of seven distinct genotypes. This indicates that, on a local scale, and even on individual birch trees, multiple distinct strains are present. Some of these strains may even represent closely related but distinct birch-associated *Taphrina* species, as will be further discussed below. There was no apparent specificity to the strains present or their diversity within the three sample types examined. *T. betulina*, like all *Taphrina* species, is dimorphic, infecting its host in the dikaryotic hyphal form in the spring and early summer, but existing as a phylloplane resident in its yeast form for most of the year. Additionally, *Taphrina* species are sensitive to environmental conditions and only infect when favourable cold and wet conditions prevail during bud break in the spring (Giosuè et al., 2000, Rossi et al., 2006, Rossi et al., 2007).Thus, infections do not occur every year and *Taphrina* species are thought to be able to survive in the phyllosphere in their yeast form indefinitely (Fonseca and Rodrigues, 2011, Mix, 1949). As such they are expected to be found even on healthy tissues on their hosts; although there is little experimental evidence supporting this. Strains of *T. betulina* with the genotype ITS-D RGR-2 were the most commonly found and exhibited a pattern where strains of this genotype were more frequently isolated from more diseased samples, making it a strong candidate for the being the primary cause of witches’ broom disease in the trees studied here. Its presence on symptom-free leaf samples supports that virulent *T. betulina* strains can also be found as phyllosphere residents on apparently healthy trees. There are only a few studies that have previously made such observations, using molecular methods *T. deformans* has been detected from healthy peach trees (Tavares et al., 2004, Mikhailova et al., 2020).

The presence of multiple strains in the same samples also suggests a diversity of *T. betulina* strains may be involved in the disease process. This is consistent with the known ability of *Taphrina* species to enter the dikaryotic state by two distinct mechanisms; either with a single cell that undergoes nuclear duplication without cell division, or by conjugation of two yeast cells. The former seems to be the dominant mechanism. The regulation of these varied sexual behaviours in *Taphrina* species are not well understood. Characterization of the *T deformans* genome indicates a MAT locus configured for primary homothallism (Almeida et al., 2015). Conjugation has only been rarely observed in a few species, *T. betulina* not among them (Mix, 1949). Further studies are required to explore the possibility of conjugation by *T. betulina* strains suggested by these findings.

Alternatively, some of the various strains present may have different specialist lifestyles. Some *Taphrina* species have been isolated from symptomless plants that were not previously known to be *Taphrina* hosts (Fonseca and Inácio, 2011, Inacio et al., 2004). These species have been found on multiple plant species, are thought to be non-pathogens specialized in phyllosphere residency, and have broader than usual carbon utilization profiles (Fonseca and Inácio, 2011, Inacio et al., 2004). Strains of *T. betulina* with four genotypes (ITC-C RGR-1, ITC-C RGR-2, ITC-D RGR-2, ITC-D RGR-3) were found only on healthy samples. Although, strains of these genotypes were also rare and further studies with deeper sampling may be required to understand their true distribution. Nonetheless, these strains raise the question of possible phyllosphere specialist *Taphrina* on birch. Examination of the carbon utilization capabilities of these strains will be necessary to address this question.

### Isolated *T. betulina* strains compared to other*Taphrina* species isolated from birch

There are many birch associated *Taphrina* species (Table 1). As is typical for all *Taphrina* species, *T. betulina* and the other birch associated *Taphrina* have a host range that is fairly wide within the genus *Betula* (Fonseca and Rodrigues, 2011, Mix, 1949). Some geographically separate species do exist; for instance *T. americana* that is pathogenic on North American *Betula* species and *T. betulina* that is pathogenic on Eurasian *Betula* species. Very few of these birch-associated *Taphrina* have been analysed with modern molecular methods and some have been lost with no viable cultures currently available.

A long running trend in research on these species has seen a reduction in their numbers through merging of conspecific lines (Mix, 1949). This calls for critical re-evaluation the many characteristics previously used to separate species, such as cell size, morphology, symptoms induced in the host, geographical location of isolation, host species, etc. In their place, reliance on molecular studies must take precedence, with which further merging of conspecific strains can be expected. This is well illustrated by the cases of *T. carnea* and *T. nana*, which were separated from *T. betulina* based on combinations of morphology, host tissues infected, and symptoms produced (Mix, 1949), but later judged to be conspecific with *T. betulina* based on identical ITS sequences and PCR finger printing patterns (Rodrigues and Fonseca, 2003). In the current work, we developed a new molecular marker named RGR that distinguishes between *T. betulina* strains. Remarkably, use of this marker and the ITS sequence based CAPS marker ITC led to the detection of strains of ITC-D RGR-2 (Table 3), a genotype that was the most commonly isolated and is identical to *T. nana* (Fig. S3). This finding suggests that the most commonly isolated *T. betulina* strain found in all sample types, but more commonly in symptomatic samples, may be *T. nana*. The species *T. nana* has been previously observed on *B. pendula* (Table 1) (Fonseca and Rodrigues, 2011, Mix, 1949) To further support this observation, we compared the cell sizes of strains with the genotype ITC-D RGR-2 (Figure 3; Table S3) to the published size ranges for *T. nana* (Table 1), which were roughly in agreement.

There is clear evidence for birch associated *Taphrina* with distinct ITS sequences; T. americana has an ITS sequence that was different at 20 nucleotide positions compared to *T. betulina*. Strains of the genotype ITC-C were also distinct from T. betulina, wich had the genotype ITC-D. Further comparisons will be required to determine if these strains represent a known or novel birch associated *Taphrina* species or strain.

## Supporting information

Christita et al Supplemental Materials

## Acknowledgements

We thank Tuomas Puukko, Airi Lamminmäki, and Leena Grönholm, for excellent technical support. This work was supported by the following grants: the Academy of Finland Center of Excellence in Primary Producers 2014-2019 (decisions #271832 and 307335); a PhD fellowship to MC from the Indonesian Fund for Education (LPDP); and a research grant to MC from the Finnish Society for Forestry Sciences. MC is a member of the University of Helsinki Doctoral Program in Plant Science (DPPS). The authors declare no conflict of interest related to this work.

## Author contributions

KO and TS conceived the research, TS and MC performed the research, KO and MC wrote the manuscript, all authors participated in data analysis, data interpretation, editing and approval of the manuscript.

## Supplemental Figure Legends

**Supplemental Figure 1. Sampled leaves**. Photos of the nine birch leaf samples (A-I) used in this study for isolation of phyllosphere yeasts. The panel labels correspond to the sample names.

**Supplemental Figure 2**. Classification of the 22 *T. betulina* strains selected for further analysis. Classification is based on the three types of host the strains were collected from; type I, symptomatic leaves from symptomatic tree; type II, asymptomatic leaves from symptomatic tree; type III, asymptomatic leaves from asymptomatic tree.

**Supplemental Figure 3**. PCR markers used to classify *Taphrina betulina* strains. **(A)** The nuclear rRNA internal transcribed spacer (ITS) TaqI cleaved amplified polymorphic sequence (CAPS) (ITC) marker based on digestion of the ITS PCR product (Containing ITS1-5S-ITS2 sequences) with the restriction endonuclease TaqI. **(B)** Schematic of new marker design. Two conserved housekeeping gene pairs were found with conserved synteny in multiple species of *Taphrina* and primers designed as depicted to allow primer binding in conserved gene regions and amplification of polymorphic intergenic regions. **(C)** Gene and primer names with their expected PCR products and expected intergenic region lengths. **(D)** Test PCR results using *T. betulina* and *T. deformans* genomic DNA as template. **(E)** Right, PCR amplification results using the gyp7-rco1 primer set with a larger collection of genomic DNA templates from *Taphrina* species. Left, gyp7-rco1 cleaved amplified polymorphic sequence (CAPS) results after PCR product digestion with the RsaI restriction endonuclease. This marker is termed the rco1 gyp7 RsaI (RGR) CAPS marker. **(F)** Alignments between *T. betulina* strain PYCC 5889 (=CBS 119536=NRRL T-726; ITS accession AF492080.1), *T*. carnea strain PYCC 5890 (=NRRL T-705; ITS accession AF492084.1), *T*. nana strain PYCC 5716 (=CBS 336.55; ITS accession AF492102) *T. robinsoniana* strain NRRL T-732 (ITS accession AF492116.1), and *T. americana* strain PYCC 5701 (ITS accession AF492078).

**Supplemental Figure 4. Colony and yeast cell morphology**. Twenty-two selected *Taphrina betulina* strains isolated from birch leaves. Organized by sample type the strain numbers were; Type I - 25, 26, 34, 85, 31; Type II - 82, 219, 58), 83, 59, 68, 69, 198, 62, 199, 151, 63, 112; Type III - 129, 11, 19, 20.

**Supplemental Figure 5. Activation of auxin dependent transcriptional response *in planta***. Response was monitored by the auxin, (indole acetic acid, IAA) -responsive DR5 promoter fused to the β-glucuronidase (*GUS*) reporter gene. Two weeks old *in vitro* grown *Arabidopsis thaliana* seedlings transgenically bearing *DR5::GUS* were treated for 24 hours with filtered supernatants of five-day-old *Taphrina betulina* cultures in YPD and YPD +0.1%tryptophan and then stained for GUS activity, which deposited an insoluble blue coloured product in tissues where the promoter was active. Uncultured media (YPD and YPD + Trp) were used as negative controls and 5 µM IAA as a positive control.

**Supplemental Table 1**. Full list of all strains isolated. Abbreviations used: nd, no data; np, no ITS PCR product

**Supplemental Table 2**. *Taphrina betulina* strain diversity from different broom morphology types.

**Supplemental Table 3**. Cell Size of 22 *T. betulina* selected strains

