## Supplementary material for "The isolation and characterization of *Taphrina betulina* and other yeasts residing in the *Betula pendula* phylloplane": Christita et al Supplemental Materials

Supplementary Fig.1

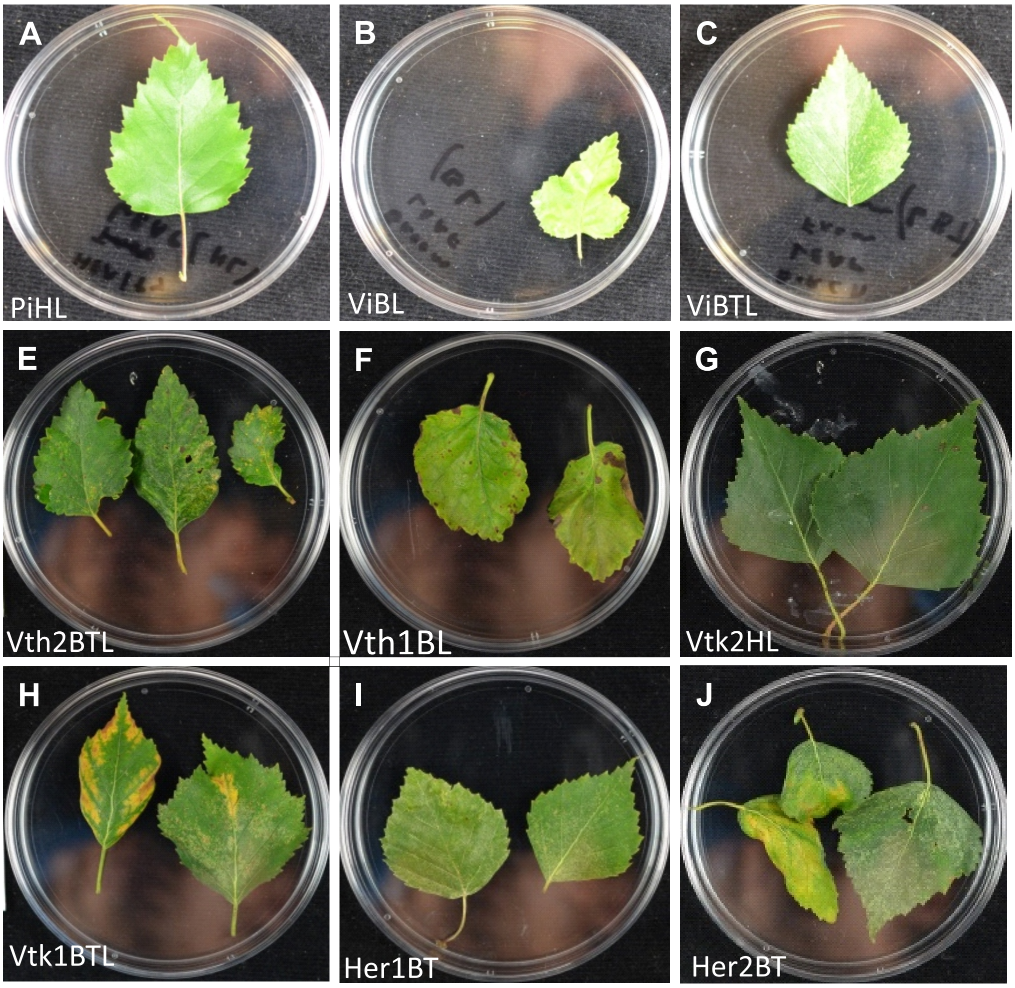


L

Supplementary Figure 2

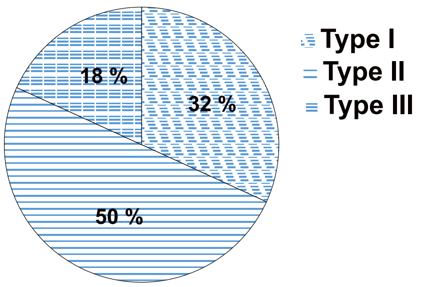

### Supplementary Figure 3

A.

| ITS TaqI CAPS marker type | Banding pattern | *Taphrina betulina type* | Closest BLAST hit |
|---|---|---|---|
| ITC-D | 248,232,132, 59, 27 | Variant I | Taphrina betulina strain NRRL T-726 100% |
| ITC-C | 323, 263,59, | Variant II | Taphrina betulina strain NRRL T-726 99% |

B

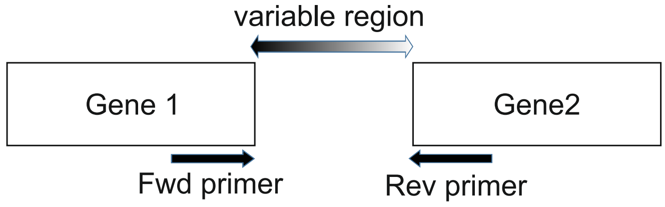


C

| Forward primer | Reverse primer | Product length (bp) | variable region (bp) |
|---|---|---|---|
| sad1 fwd | rax1 rev | 1250 | 587 |
| gyp7 fwd | rco1 rev | 1200 | 497 |

D

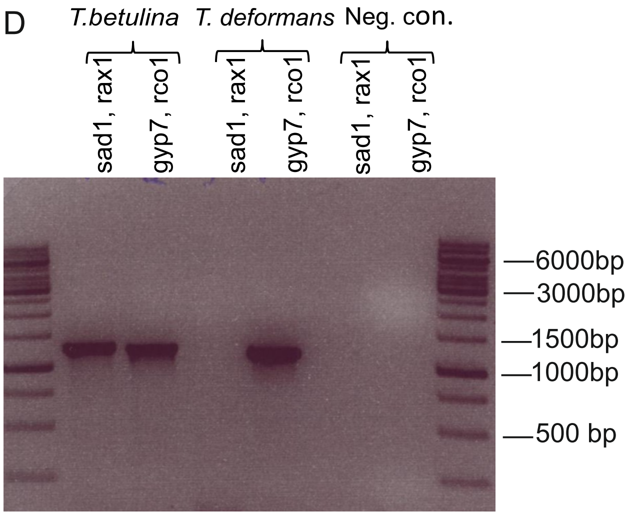


E

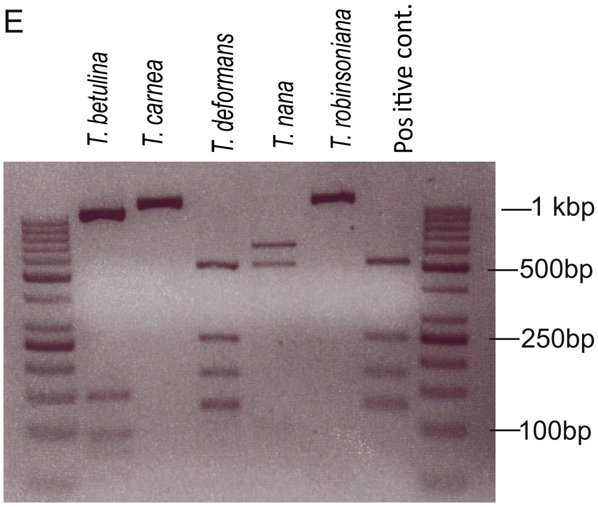


| Species | Strain | RGR PCR | RGR type |
|---|---|---|---|
| *T. betulina* | PYCC 5889 | + | RGR-3 |
| *T. carnea* | PYCC 5890 | + | RGR-1 |
| *T. deformans* | PYCC 5710 | + | - |
| *T. americana* | PYCC 5071 | | RGR-0 |
| *T. nana* | PYCC 5716 | + | RGR-2 |
| *T. robinsoniana* | T-732 | + | RGR-1 |

F

```
T robinsoniana      AAGGATCATTAATGAAGTCTGGGCTTCGGCCCTCTCTCTTCTACACACTTGTGAATCTAC     60
T americana         AAGGATCATTAATGAAGTCTGGGCTTCGGCCCTCTCTCTTCTACACACTTGTGAACTTAC     60
T nana              AAGGATCATTAATGAAGTCTGGGCTCCGGCCCTCTCTCTTCTACACACTTGTGAACTTAC     60
T carnea            AAGGATCATTAATGAAGTCTGGGCTCCGGCCCTCTCTCTTCTACACACTTGTGAACTTAC     60
T betulina          AAGGATCATTAATGAAGTCTGGGCTCCGGCCCTCTCTCTTCTACACACTTGTGAACTTAC     60
                    ************************* *****************************  ***
                                                      ITS1

T robinsoniana      ACTGTTGCTTTGGCAGGTAGCCGGACGGACGTGAGTCTGCACGGCGAGGTCGAGAGACGC     120
T americana         ACCGTTGCTTTGGCAGGTTTCCGGAGGGGCGAAAGCTTCGAAGGTCAGGTCGAAAGGCGC     120
T nana              ACTGTTGCTTTGGCAGGTTTCCGGACGGGCGAAAGCTCTGAAGGTCAGGTCGAAAGGCGC     120
T carnea            ACTGTTGCTTTGGCAGGTTTCCGGACGGGCGAAAGCTCTGAAGGTCAGGTCGAAAGGCGC     120
T betulina          ACTGTTGCTTTGGCAGGTTTCCGGACGGGCGAAAGCTCTGAAGGTCAGGTCGAAAGGCGC     120
                    ** ***************  ***** ** **  **     * **  ******* ** ***
                                              ITS1

T robinsoniana      CTGCCAAGGACATTTATCCACCCTTTTTCAATAGTCTGATTATTGTTTTAAACAAATTAA     180
T americana         CTGCCAAGGACATTTATCCACCCTTTTTATATCGTCTGATTTTTGTTTTAAACAAATTAT     180
T nana              CTGCCAAGGACATTTACCCACCCTTTTTATATTGTCTGATTTTTGTTTTAAACAAATTAT     180
T carnea            CTGCCAAGGACATTTACCCACCCTTTTTATATTGTCTGATTTTTGTTTTAAACAAATTAT     180
T betulina          CTGCCAAGGACATTTACCCACCCTTTTTATATTGTCTGATTTTTGTTTTAAACAAATTAT     180
                    **************** ***********  ** ******** ***************** 
                                              ITS1

T robinsoniana      AATAAAACTTTCAACAATGGATCTCTTGGCTCTGGCATCGATGAAGAACGCAGCGAAATG     240
T americana         AATAAAACTTTCAACAATGGATCTCTTGGCTCTGGCATCGATGAAGAACGCAGCGAAATG     240
T nana              AATAAAACTTTCAACAATGGATCTCTTGGCTCTGGCATCGATGAAGAACGCAGCGAAATG     240
T carnea            AATAAAACTTTCAACAATGGATCTCTTGGCTCTGGCATCGATGAAGAACGCAGCGAAATG     240
T betulina          AATAAAACTTTCAACAATGGATCTCTTGGCTCTGGCATCGATGAAGAACGCAGCGAAATG     240
                    ************************************************************
                                            5S rRNA

T robinsoniana      CGATAAGTAATGTGAATTGCAGAATTCAGTGAATCATCGAATCTTTGAACGCACATTGCG     300
T americana         CGATAAGTAATGTGAATTGCAGAATTCAGTGAATCATCGAATCTTTGAACGCACATTGCG     300
T nana              CGATAAGTAATGTGAATTGCAGAATTCAGTGAATCATCGAATCTTTGAACGCACATTGCG     300
T carnea            CGATAAGTAATGTGAATTGCAGAATTCAGTGAATCATCGAATCTTTGAACGCACATTGCG     300
T betulina          CGATAAGTAATGTGAATTGCAGAATTCAGTGAATCATCGAATCTTTGAACGCACATTGCG     300
                    ************************************************************
                                            5S rRNA
```

```
T robinsoniana      CCCTCTGGTATTCCGGAGGGCATGCCTGTTTGAGTGTCATTAATCTCTCACAAAGACCTT    360
T americana         CCCTCTGGTATTCCGGAGGGCATGCCTGTTTGAGTGTCATTAATCTCTCACAAAAC-CTT    359
T nana              CCCTCTGGTATTCCGGAGGGCATGCCTGTTTGAGTGTCATTAACTTCTCACAAAAC-CTT    359
T carnea            CCCTCTGGTATTCCGGAGGGCATGCCTGTTTGAGTGTCATTAACTTCTCACAAAAC-CTT    359
T betulina          CCCTCTGGTATTCCGGAGGGCATGCCTGTTTGAGTGTCATTAACTTCTCACAAAAC-CTT    359
                    *******************************************  *********   ***
                    [              5S rRNA                      ][    ITS2     ]

T robinsoniana      TTGGTCTCTGGTGAAGTTGGGATCTGCGTACCTTTGTGGAACGCTGTCCCCAAATAGATT    420
T americana         TTGGTTTCTGTCGATGTTGGGAGCTGCGACCCCTCGTGGGACGCTCTCCTTAAATGTATT    419
T nana              T-----------------------------------------------------------    360
T carnea            TTGGTTTCTGTTGATGTTGGGAACTGCGACCCCTCGTGGGACGCTTTCCTCAAATGCATT    419
T betulina          TTGGTTTCTGTTGATGTTGGGAACTGCGACCCCTCGTGGGACGCTTTCCTCAAATGCATT    419
                    *
                    [                          ITS2                            ]

T robinsoniana      GGTGCGGCCCGCCGCCC-GGTACACAACGTTCTAGGTTCGTCCAACTCGTTGATCAACCG    479
T americana         GGTGCGGCCCGCCGCCCGGTTACACAACGTTCTAGGTTCGTCCAACTCGTTGCTCAGCCG    479
T nana              ------------------------------------------------------------    360
T carnea            GGTGCGGCCCGCCGCCCGGTAACACAACGTTCTAGGTTCGTCCAACTCGTTGCTCGC-CG    478
T betulina          GGTGCGGCCCGCCGCCCGGTAACACAACGTTCTAGGTTCGTCCAACTCGTTGCTCGC-CG    478

                    [                          ITS2                            ]

T robinsoniana      GTGCATATATGGTGCTGCACCTAAAGCCCGCCTTGTGCCTTGTGCTCTTGGCTCTTTT-A    538
T americana         GTGCAATCTTGGTGCTGCACCTAAAGCCCGCCTCGTGCCTTGTGCTCTTGGCTAACTTCA    539
T nana              ------------------------------------------------------------    360
T carnea            GTGCAATCTTGGTGCTGCACCTTAAGCCCGCCCTGTGCCTTGTGCACTTGGCTAACTTCA    538
T betulina          GTGCAATCTTGGTGCTGCACCTTAAGCCCGCCCTGTGCCTTGTGCACTTGGCTAACTTCA    538

                    [                          ITS2                            ]

T robinsoniana      TCTATTGACCTCAGATCAGGTAGGAATACGCGCTGAACTTAAGC    582
T americana         T-TATTGACCTCAGATCAGGTAGGAATACGCGCTGAACTTAAGC    582
T nana              --------------------------------------------    360
T carnea            TTTATTGACCTCAGATCAGGTAGGAATACGCGCTGAACTTAAGC    582
T betulina          TTTATTGACCTCAGATCAGGTAGGAATACGCGCTGAACTTAAGC    582
```

Supplementary Fig 4.

Type I (Symptomatic leaf - symptomatic tree)

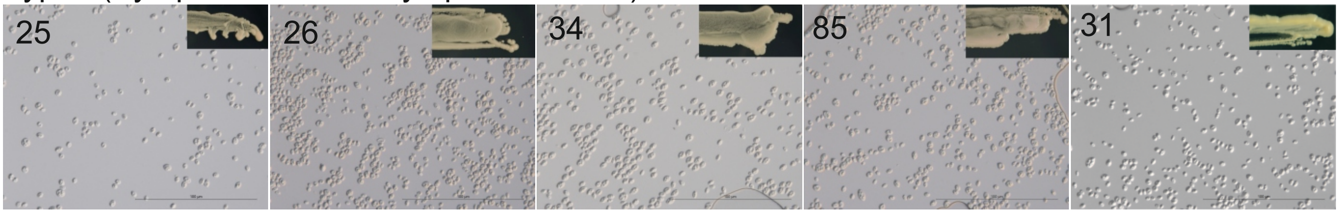


Type II (Asymptomatic leaf - symptomatic tree)

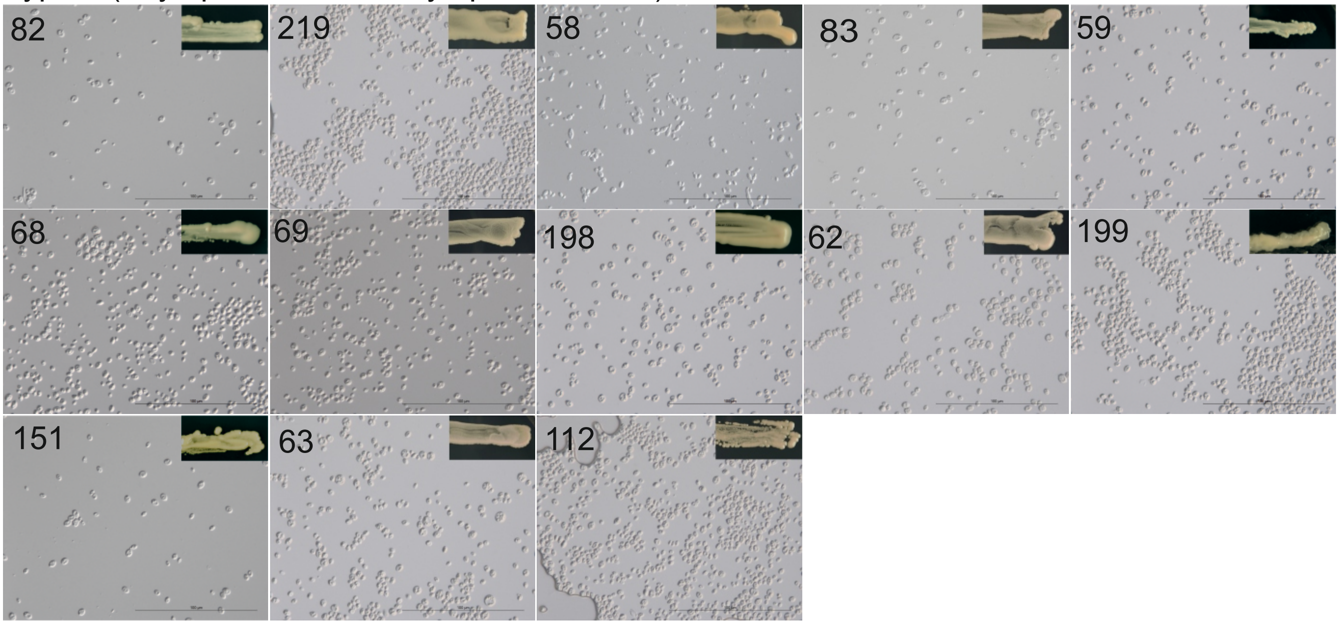


Type III (Asymptomatic leaf - asymptomatic tree)

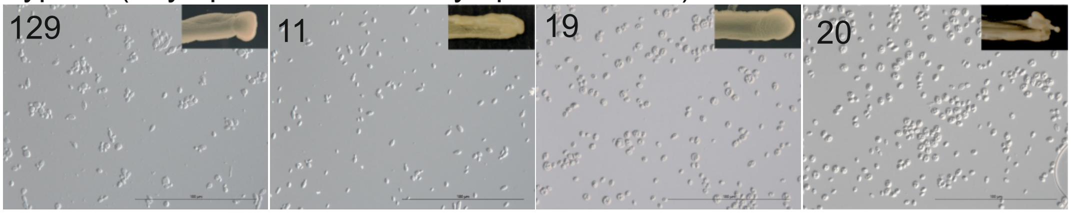

### Supplementary Figure 5

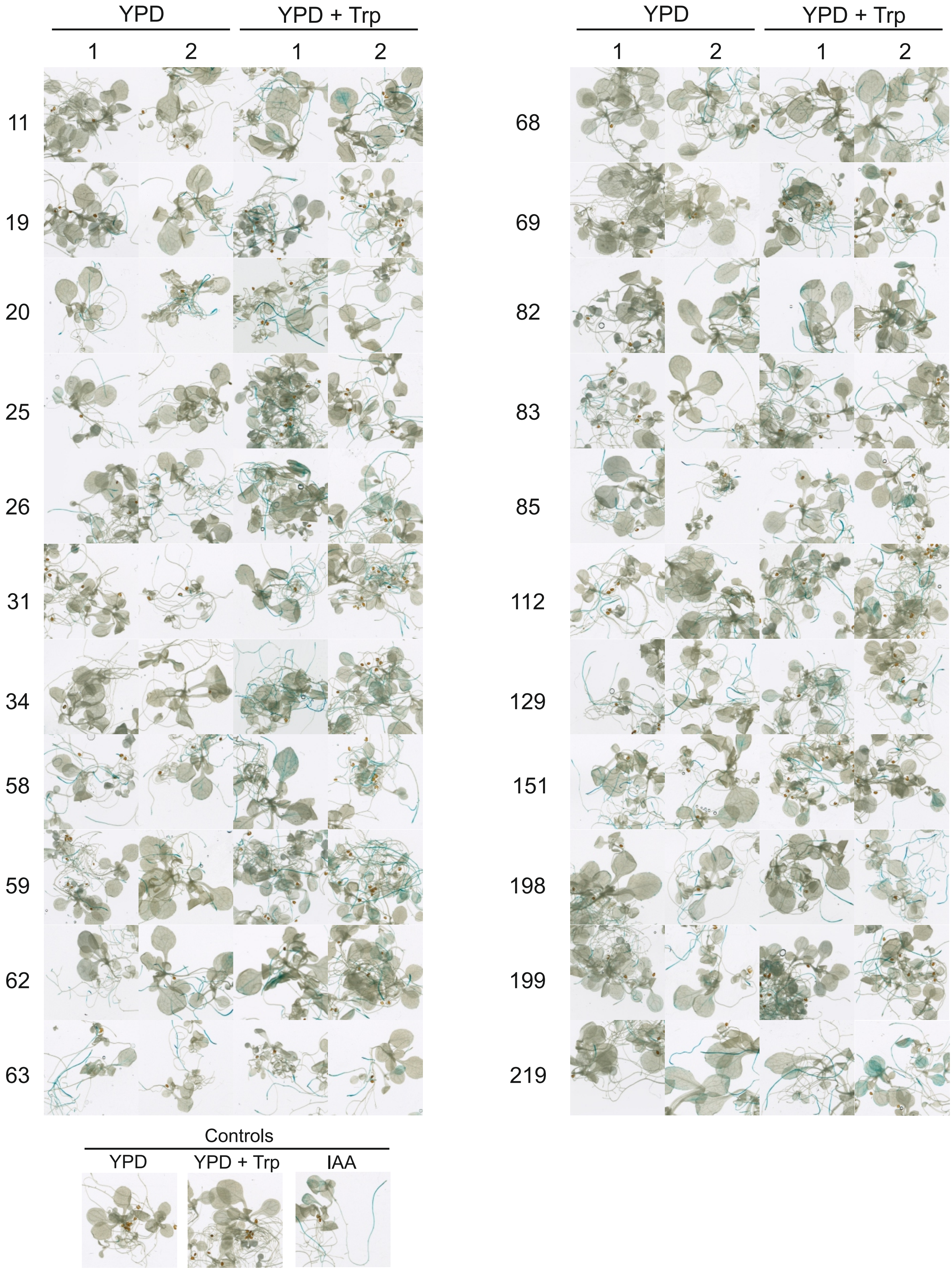

**Supplementary Table 1: Full list of all strains isolated.** [a] Samples are defined in Table 1. [B]Strain growth was measured by drop inoculations of serial dilutions on 0.2 X PDA plates and incubation at the indicated temperature for four days then were visually assessed and characterized as noted: +++, strong growth; ++, intermediate growth; + weak growth. [c]Nuclear rRNA internal transcribed spacer (ITS) PCR products were used as cleaved amplified polymorphic sequence (CAPS) marker, called ITS Taq I CAPS (ITC), by digestion with the restriction endonuclease Taq I. ITC types represent distinct banding patters (ITC-A to ITC-T). [d]RGR (Rco1 Gyg7 Rsa1) is a CAPS marker that amplifies the polymorphic intergenic region between the *Taphrina betulina* Gyg7 and Rco1 genes and is digested with the restriction nuclease RsaI. It is used to differential between *T. betulina* strains and identifies four different banding patters RGR-0, no PCR product, and RGR-1 to RGR-3, which represent banding pattern variants I-III, respectively. See Supplemental Figure 3 for further details on the design of CAPS markers. Abbreviations used: nt, not tested; nd, no data; np, no ITS PCR product; ui, unidentified.

| Strain | Location | Tree phenotype | Isolation source | Sample[a] | Growth 30°C[b] | Growth 21°C[b] | ITC type[c] | RGR type[d] |
|---|---|---|---|---|---|---|---|---|
| **1** | Pihlajisto | Small healthy | Healthy appearing leaf | A | nd | nd | np | nt |
| **2** | Pihlajisto | Small healthy | Healthy appearing leaf | A | - | +++ | ITC-E | nt |
| **3** | Pihlajisto | Small healthy | Healthy appearing leaf | A | - | +++ | ITC-E | nt |
| **4** | Pihlajisto | Small healthy | Healthy appearing leaf | A | - | +++ | ITC-E | nt |
| **5** | Pihlajisto | Small healthy | Healthy appearing leaf | A | - | +++ | ITC-E | nt |
| **6** | Pihlajisto | Small healthy | Healthy appearing leaf | A | - | +++ | ITC-E | nt |
| **7** | Pihlajisto | Small healthy | Healthy appearing leaf | A | - | +++ | ITC-F | nt |
| **8** | Pihlajisto | Small healthy | Healthy appearing leaf | A | - | +++ | ITC-E | nt |
| **9** | Pihlajisto | Small healthy | Healthy appearing leaf | A | - | + | ITC-E | nt |
| **10** | Pihlajisto | Small healthy | Healthy appearing leaf | A | - | + | ITC-E | nt |
| **11** | Pihlajisto | Small healthy | Healthy appearing leaf | A | - | + | ITC-C | RGR-1 |
| **12** | Pihlajisto | Small healthy | Healthy appearing leaf | A | - | +++ | ITC-E | nt |
| **13** | Pihlajisto | Small healthy | Healthy appearing leaf | A | - | + | ITC-E | nt |
| **14** | Pihlajisto | Small healthy | Healthy appearing leaf | A | - | +++ | ITC-E | nt |
| **15** | Pihlajisto | Small healthy | Healthy appearing leaf | A | - | +++ | ITC-E | nt |
| **16** | Pihlajisto | Small healthy | Healthy appearing leaf | A | - | +++ | ITC-E | nt |
| **17** | Pihlajisto | Small healthy | Healthy appearing leaf | A | - | +++ | ITC-E | nt |
| **18** | Pihlajisto | Small healthy | Healthy appearing leaf | A | - | + | ITC-E | nt |
| **19** | Pihlajisto | Small healthy | Healthy appearing leaf | A | - | +++ | ITC-D | RGR-0 |
| **20** | Pihlajisto | Small healthy | Healthy appearing leaf | A | - | +++ | ITC-D | RGR-2 |
| **21** | Pihlajisto | Small healthy | Healthy appearing leaf | A | + | +++ | ITC-D | RGR-2 |
| **22** | Pihlajisto | Small healthy | Healthy appearing leaf | A | - | +++ | ITC-D | RGR-2 |
| **23** | Pihlajisto | Small healthy | Healthy appearing leaf | A | - | +++ | ITC-F | nt |
| **24** | Pihlajisto | Small healthy | Healthy appearing leaf | A | - | +++ | ITC-E | nt |
| **25** | Viikinkaari | Large birch with elongated brooms | Broom leaf small and deformed | B | - | +++ | ITC-D | RGR-2 |
| **26** | Viikinkaari | Large birch with elongated brooms | Broom leaf small and deformed | B | - | +++ | ITC-D | RGR-2 |
| **27** | Viikinkaari | Large birch with elongated brooms | Broom leaf small and deformed | B | + | + | ITC-D | RGR-0 |
| **28** | Viikinkaari | Large birch with elongated brooms | Broom leaf small and deformed | B | + | +++ | np | nt |
| **29** | Viikinkaari | Large birch with elongated brooms | Broom leaf small and deformed | B | - | +++ | ITC-H | nt |
| **30** | Viikinkaari | Large birch with elongated brooms | Broom leaf small and deformed | B | - | +++ | ITC-D | RGR-2 |
| **31** | Viikinkaari | Large birch with elongated brooms | Broom leaf small and deformed | B | - | +++ | ITC-D | RGR-0 |
| **32** | Viikinkaari | Large birch with elongated brooms | Broom leaf small and deformed | B | - | ++ | ITC-D | RGR-2 |
| **33** | Viikinkaari | Large birch with elongated brooms | Broom leaf small and deformed | B | - | +++ | ITC-D | RGR-2 |
| **34** | Viikinkaari | Large birch with elongated brooms | Broom leaf small and deformed | B | - | + | ITC-D | RGR-2 |
| **35** | Viikinkaari | Large birch with elongated brooms | Broom leaf small and deformed | B | + | +++ | ITC-F | nt |
| **36** | Viikinkaari | Large birch with elongated brooms | Broom leaf small and deformed | B | - | +++ | ITC-D | RGR-2 |

| | | | | | | | | |
|---|---|---|---|---|---|---|---|---|
| **37** | Viikinkaari | Large birch with elongated brooms | Broom leaf small and deformed | B | - | +++ | ITC-D | RGR-2 |
| **38** | Viikinkaari | Large birch with elongated brooms | Broom leaf small and deformed | B | - | +++ | ITC-J | nt |
| **39** | Viikinkaari | Large birch with elongated brooms | Broom leaf small and deformed | B | - | - | ITC-D | RGR-0 |
| **40** | Viikinkaari | Large birch with elongated brooms | Broom leaf small and deformed | B | - | +++ | ITC-N | nt |
| **41** | Viikinkaari | Large birch with elongated brooms | Broom leaf small and deformed | B | + | +++ | ITC-O | nt |
| **42** | Viikinkaari | Large birch with elongated brooms | Broom leaf small and deformed | B | + | +++ | np | nt |
| **43** | Viikinkaari | Large birch with elongated brooms | Broom leaf small and deformed | B | - | - | ITC-D | RGR-0 |
| **44** | Viikinkaari | Large birch with elongated brooms | Broom leaf small and deformed | B | - | +++ | ITC-D | RGR-2 |
| **45** | Viikinkaari | Large birch with elongated brooms | Broom leaf small and deformed | B | + | +++ | np | nt |
| **46** | Viikinkaari | Large birch with elongated brooms | Broom leaf small and deformed | B | - | - | ITC-P | nt |
| **47** | Viikinkaari | Large birch with elongated brooms | Broom leaf small and deformed | B | - | - | ITC-P | nt |
| **48** | Viikinkaari | Large birch with elongated brooms | Broom leaf small and deformed | B | - | - | ITC-Q | nt |
| **49** | Viikinkaari | Large birch with elongated brooms | Broom leaf small and deformed | B | - | - | ITC-D | RGR-2 |
| **50** | Viikinkaari | Large birch with elongated brooms | Broom leaf small and deformed | B | - | +++ | ITC-D | RGR-2 |
| **51** | Viikinkaari | Large birch with elongated brooms | Broom leaf small and deformed | B | - | - | ITC-D | RGR-0 |
| **52** | Viikinkaari | Large birch with elongated brooms | Broom leaf small and deformed | B | + | +++ | np | nt |
| **53** | Viikinkaari | Large birch with elongated brooms | Broom leaf small and deformed | B | - | ++ | ITC-D | RGR-2 |
| **54** | Viikinkaari | Large birch with elongated brooms | Broom tree leaf with gray small spot | C | + | +++ | ITC-F | nt |
| **55** | Viikinkaari | Large birch with elongated brooms | Broom tree leaf with gray small spot | C | + | + | ITC-F | nt |
| **56** | Viikinkaari | Large birch with elongated brooms | Broom tree leaf with gray small spot | C | + | +++ | ITC-R | nt |
| **57** | Viikinkaari | Large birch with elongated brooms | Broom tree leaf with gray small spot | C | - | +++ | ITC-D | RGR-2 |
| **58** | Viikinkaari | Large birch with elongated brooms | Broom tree leaf with gray small spot | C | - | +++ | ITC-D | RGR-3 |
| **59** | Viikinkaari | Large birch with elongated brooms | Broom tree leaf with gray small spot | C | - | +++ | ITC-D | RGR-2 |
| **60** | Viikinkaari | Large birch with elongated brooms | Broom tree leaf with gray small spot | C | - | +++ | ITC-I | nt |
| **61** | Viikinkaari | Large birch with elongated brooms | Broom tree leaf with gray small spot | C | - | ++ | ITC-R | nt |
| **62** | Viikinkaari | Large birch with elongated brooms | Broom tree leaf with gray small spot | C | - | +++ | ITC-D | RGR-2 |
| **63** | Viikinkaari | Large birch with elongated brooms | Broom tree leaf with gray small spot | C | - | + | ITC-C | RGR-2 |
| **64** | Viikinkaari | Large birch with elongated brooms | Broom tree leaf with gray small spot | C | - | + | ITC-D | RGR-0 |
| **65** | Viikinkaari | Large birch with elongated brooms | Broom tree leaf with gray small spot | C | + | +++ | ITC-R | nt |
| **66** | Viikinkaari | Large birch with elongated brooms | Broom tree leaf with gray small spot | C | + | +++ | ITC-R | nt |
| **67** | Viikinkaari | Large birch with elongated brooms | Broom tree leaf with gray small spot | C | + | +++ | ITC-D | RGR-2 |
| **68** | Viikinkaari | Large birch with elongated brooms | Broom tree leaf with gray small spot | C | + | +++ | ITC-C | RGR-2 |
| **69** | Viikinkaari | Large birch with elongated brooms | Broom tree leaf with gray small spot | C | + | +++ | ITC-D | RGR-2 |
| **70** | Viikinkaari | Large birch with elongated brooms | Broom tree leaf with gray small spot | C | + | +++ | np | nt |
| **71** | Viikinkaari | Large birch with elongated brooms | Broom tree leaf with gray small spot | C | + | +++ | np | nt |
| **72** | Viikinkaari | Large birch with elongated brooms | Broom tree leaf with gray small spot | C | - | +++ | ITC-D | RGR-2 |
| **73** | Viikinkaari | Large birch with elongated brooms | Broom tree leaf with gray small spot | C | + | +++ | ITC-F | nt |
| **74** | Viikinkaari | Large birch with elongated brooms | Broom tree leaf with gray small spot | C | - | +++ | ITC-D | RGR-2 |
| **75** | Viikinkaari | Large birch with elongated brooms | Broom tree leaf with gray small spot | C | - | +++ | ITC-D | RGR-2 |
| **76** | Viikinkaari | Large birch with elongated brooms | Broom tree leaf with gray small spot | C | + | +++ | ITC-J | nt |
| **77** | Pihlajisto | Small healthy | Healthy appearing leaf | A | - | +++ | np | nt |
| **78** | Pihlajisto | Small healthy | Healthy appearing leaf | A | + | +++ | ITC-I | nt |
| **79** | Pihlajisto | Small healthy | Healthy appearing leaf | A | + | +++ | ITC-I | nt |
| **80** | Pihlajisto | Small healthy | Healthy appearing leaf | A | + | +++ | ITC-E | nt |
| **81** | Pihlajisto | Small healthy | Healthy appearing leaf | A | + | +++ | ITC-E | nt |
| **82** | Viikinkaari | Large birch with elongated brooms | Broom tree leaf with gray small spot | C | - | +++ | ITC-D | RGR-1 |
| **83** | Viikinkaari | Large birch with elongated brooms | Broom tree leaf with gray small spot | C | - | +++ | ITC-D | RGR-0 |

| | | | | | | | | |
|---|---|---|---|---|---|---|---|---|
| **84** | Viikinkaari | Large birch with elongated brooms | Broom leaf small and deformed | B | + | +++ | ITC-J | nt |
| **85** | Viikinkaari | Large birch with elongated brooms | Broom leaf small and deformed | B | - | +++ | ITC-C | RGR-0 |
| **86** | Vartioharju (VTH) | Large birch hanging phenotype elongated brooms, heavily diseased (VTHS1) | Broom tree leaf with gray small spots and yellow spot and deficient growth (BTL) | D | - | +++ | ITC-A | nt |
| **87** | Vartioharju (VTH) | Large birch hanging phenotyp) with elongated brooms, heavily diseased (VTHS1) | Broom tree leaf with gray small spots and yellow spot and deficient growth (BTL) | D | + | +++ | ITC-F | nt |
| **88** | Vartioharju (VTH) | Large birch hanging phenotype elongated brooms, heavily diseased (VTHS1) | Broom tree leaf with gray small spots and yellow spot and deficient growth (BTL) | D | + | +++ | ITC-F | nt |
| **89** | Vartioharju (VTH) | Large birchhanging phenotype elongated brooms, heavily diseased (VTHS1) | Broom tree leaf with gray small spots and yellow spot and deficient growth (BTL) | D | + | +++ | ITC-F | nt |
| **90** | Vartioharju (VTH) | Large birch hanging phenotype with elongated brooms, heavily diseased (VTHS1) | Broom tree leaf with gray small spots and yellow spot and deficient growth (BTL) | D | + | +++ | ITC-F | nt |
| **91** | Vartioharju (VTH) | Large birch hanging phenotype elongated brooms, heavily diseased (VTHS1) | Broom tree leaf with gray small spots and yellow spot and deficient growth (BTL) | D | + | +++ | ITC-F | nt |
| **92** | Vartioharju (VTH) | Large birch hanging phenotype with elongated brooms, heavily diseased (VTHS1) | Broom tree leaf with gray small spots and yellow spot and deficient growth (BTL) | D | + | +++ | ITC-F | nt |
| **93** | Vartioharju (VTH) | Large birch(hanging phenotype elongated brooms, heavily diseased (VTHS1) | Broom tree leaf with gray small spots and yellow spot and deficient growth (BTL) | D | + | +++ | ITC-A | nt |
| **94** | Vartioharju (VTH) | Large birch hanging phenotype elongated brooms, heavily diseased (VTHS1) | Broom tree leaf with gray small spots and yellow spot and deficient growth (BTL) | D | + | +++ | ITC-A | nt |
| **95** | Vartioharju (VTH) | Large birch hanging phenotype elongated brooms, heavily diseased (VTHS1) | Broom tree leaf with gray small spots and yellow spot and deficient growth (BTL) | D | + | +++ | ITC-F | nt |
| **96** | Vartioharju (VTH) | Large birch hanging phenotype elongated brooms, heavily diseased (VTHS1) | Broom tree leaf with gray small spots and yellow spot and deficient growth (BTL) | D | + | +++ | ITC-F | nt |
| **97** | Vartioharju (VTH) | Large birch hanging phenotype elongated brooms, heavily diseased (VTHS1) | Broom tree leaf with gray small spots and yellow spot and deficient growth (BTL) | D | - | +++ | ITC-F | nt |
| **98** | Vartioharju (VTH) | Large birch(hanging phenotype) with elongated brooms, heavily diseased (VTHS1) | Broom tree leaf with gray small spots and yellow spot and deficient growth (BTL) | D | - | + | ITC-F | nt |
| **99** | Vartioharju (VTH) | Large birch hanging phenotype elongated brooms, heavily diseased (VTHS1) | Broom tree leaf with gray small spots and yellow spot and deficient growth (BTL) | D | +++ | +++ | np | nt |
| **100** | Vartioharju (VTH) | Large birch hanging phenotype elongated brooms, heavily diseased (VTHS1) | Broom tree leaf with gray small spots and yellow spot and deficient growth (BTL) | D | +++ | +++ | np | nt |
| **101** | Vartioharju (VTH) | Large birch hanging phenotype elongated brooms, heavily diseased (VTHS1) | Broom tree leaf with gray small spots and yellow spot and deficient growth (BTL) | D | +++ | +++ | np | nt |
| **102** | Vartioharju (VTH) | Large birch hanging phenotype elongated brooms, heavily diseased (VTHS1) | Broom tree leaf with gray small spots and yellow spot and deficient growth (BTL) | D | +++ | +++ | np | nt |
| **103** | Vartioharju (VTH) | Large birch hanging phenotype elongated brooms, heavily diseased (VTHS1) | Broom tree leaf with gray small spots and yellow spot and deficient growth (BTL) | D | +++ | +++ | np | nt |
| **104** | Vartioharju (VTH) | Large birch hanging phenotype elongated brooms, heavily diseased (VTHS1) | Broom leaf gray spots and yellow spot and deficient growth | E | +++ | +++ | np | nt |
| **105** | Vartioharju (VTH) | Large birch hanging phenotype elongated brooms, heavily diseased (VTHS1) | Broom leaf gray spots and yellow spot and deficient growth | E | +++ | +++ | np | nt |
| **106** | Vartioharju (VTH) | Large birch hanging phenotype elongated brooms, heavily diseased (VTHS1) | Broom leaf gray spots and yellow spot and deficient growth | E | - | +++ | ITC-L | nt |
| **107** | Vartioharju (VTH) | Large birch hanging phenotype | Broom leaf gray spots and yellow spot and deficient growth | E | ++ | ++ | np | nt |

| | | | | | | | | |
|---|---|---|---|---|---|---|---|---|
| | | elongated brooms, heavily diseased (VTHS1) | | | | | | |
| **108** | Vartioharju (VTH) | Large birch hanging phenotype elongated brooms, heavily diseased (VTHS1) | Broom leaf gray spots and yellow spot and deficient growth | E | ++ | + | np | nt |
| **109** | Vartioharju (VTH) | Large birch hanging phenotype with elongated brooms, heavily diseased (VTHS1) | Broom leaf gray spots and yellow spot and deficient growth | E | - | +++ | ITC-L | nt |
| **110** | Vartioharju (VTH) | Large birch hanging phenotype elongated brooms, heavily diseased (VTHS1) | Broom leaf gray spots and yellow spot and deficient growth | E | ++ | +++ | np | nt |
| **111** | Vartioharju (VTH) | Large birch hanging phenotype elongated brooms, heavily diseased (VTHS1) | Broom leaf gray spots and yellow spot and deficient growth | E | - | +++ | np | nt |
| **112** | Vartioharju (VTH) | Large birch hanging phenotype elongated brooms, heavily diseased (VTHS1) | Broom leaf gray spots and yellow spot and deficient growth | E | + | +++ | ITC-D | RGR-0 |
| **113** | Vartioharju (VTH) | Large birch hanging phenotype elongated brooms, heavily diseased (VTHS1) | Broom leaf gray spots and yellow spot and deficient growth | E | ++ | +++ | np | nt |
| **114** | Vartioharju (VTH) | Large birch hanging phenotype elongated brooms, heavily diseased (VTHS1) | Broom leaf gray spots and yellow spot and deficient growth | E | ++ | +++ | np | nt |
| **115** | Vartioharju (VTH) | Large birch hanging phenotype elongated brooms, heavily diseased (VTHS1) | Broom leaf gray spots and yellow spot and deficient growth | E | ++ | +++ | np | nt |
| **116** | Vartioharju (VTH) | Large birch hanging phenotype elongated brooms, heavily diseased (VTHS1) | Broom leaf gray spots and yellow spot and deficient growth | E | ++ | +++ | np | nt |
| **117** | Vartioharju (VTH) | Large birch hanging phenotype elongated brooms, heavily diseased (VTHS1) | Broom leaf gray spots and yellow spot and deficient growth | E | ++ | +++ | ITC-M | nt |
| **118** | Vartioharju (VTH) | Large birch hanging phenotype elongated brooms, heavily diseased (VTHS1) | Broom leaf gray spots and yellow spot and deficient growth | E | - | + | ITC-M | nt |
| **119** | Vartioharju (VTH) | Large birch hanging phenotype elongated brooms, heavily diseased (VTHS1) | Broom leaf gray spots and yellow spot and deficient growth | E | ++ | +++ | np | nt |
| **120** | Vartioharju (VTH) | Large birch hanging phenotype elongated brooms, heavily diseased (VTHS1) | Broom leaf gray spots and yellow spot and deficient growth | E | ++ | +++ | ITC-D | RGR-0 |
| **121** | Vartioharju (VTH) | Large birch hanging phenotype elongated brooms, heavily diseased (VTHS1) | Broom leaf gray spots and yellow spot and deficient growth | E | - | +++ | ITC-F | nt |
| **122** | Vartioharju (VTH) | Large birch hanging phenotype elongated brooms, heavily diseased (VTHS1) | Broom leaf gray spots and yellow spot and deficient growth | E | - | - | np | nt |
| **123** | Vartioharju (VTH) | Large birch hanging phenotype elongated brooms, heavily diseased (VTHS1) | Broom leaf gray spots and yellow spot and deficient growth | E | ++ | +++ | np | nt |
| **124** | Vartioharju (VTH) | Large birch hanging phenotype elongated brooms, heavily diseased (VTHS1) | Broom leaf gray spots and yellow spot and deficient growth | E | ++ | +++ | np | nt |
| **125** | Vartioharju (VTH) | Large birch hanging phenotype elongated brooms, heavily diseased (VTHS1) | Broom leaf gray spots and yellow spot and deficient growth | E | ++ | +++ | ITC-M | nt |
| **126** | Vartioharju (VTH) | Large birch hanging phenotype elongated brooms, heavily diseased (VTHS1) | Broom leaf gray spots and yellow spot and deficient growth | E | ++ | +++ | ITC-M | nt |
| **127** | Vartiokylä (VTKR2) | Large healthy birch (near heavily diseased tree) | Healthy appearing leaf | F | - | ++ | ITC-A | nt |
| **128** | Vartiokylä (VTKR2) | Large healthy birch (near heavily diseased tree) | Healthy appearing leaf | F | - | + | ITC-E | nt |
| **129** | Vartiokylä (VTKR2) | Large healthy birch (near heavily diseased tree) | Healthy appearing leaf | F | - | +++ | ITC-D | RGR-2 |

| | | | | | | | | |
|---|---|---|---|---|---|---|---|---|
| **130** | Vartiokylä (VTKR2) | Large healthy birch (near heavily diseased tree) | Healthy appearing leaf | F | - | + | ITC-E | nt |
| **131** | Vartiokylä (VTKR2) | Large healthy birch (near heavily diseased tree) | Healthy appearing leaf | F | - | ++ | ITC-A | nt |
| **132** | Vartiokylä (VTKR2) | Large healthy birch (near heavily diseased tree) | Healthy appearing leaf | F | - | ++ | ITC-F | nt |
| **133** | Vartiokylä (VTKR2) | Large healthy birch (near heavily diseased tree) | Healthy appearing leaf | F | - | +++ | ITC-A | nt |
| **134** | Vartiokylä (VTKR2) | Large healthy birch (near heavily diseased tree) | Healthy appearing leaf | F | - | + | ITC-A | nt |
| **135** | Vartiokylä (VTKR2) | Large healthy birch (near heavily diseased tree) | Healthy appearing leaf | F | - | +++ | ITC-A | nt |
| **136** | Vartiokylä (VTKR2) | Large healthy birch (near heavily diseased tree) | Healthy appearing leaf | F | - | +++ | ITC-F | nt |
| **137** | Vartiokylä (VTKR2) | Large healthy birch (near heavily diseased tree) | Healthy appearing leaf | F | - | +++ | ITC-A | nt |
| **138** | Vartiokylä (VTKR2) | Large healthy birch (near heavily diseased tree) | Healthy appearing leaf | F | - | +++ | ITC-A | nt |
| **139** | Vartiokylä (VTKR2) | Large healthy birch (near heavily diseased tree) | Healthy appearing leaf | F | - | +++ | ITC-A | nt |
| **140** | Vartiokylä (VTKR2) | Large healthy birch (near heavily diseased tree) | Healthy appearing leaf | F | - | +++ | ITC-A | nt |
| **141** | Vartiokylä (VTKR2) | Large healthy birch (near heavily diseased tree) | Healthy appearing leaf | F | - | +++ | ITC-A | nt |
| **142** | Vartiokylä (VTKR2) | Large healthy birch (near heavily diseased tree) | Healthy appearing leaf | F | - | +++ | ITC-A | nt |
| **143** | Vartiokylä (VTKR2) | Large healthy birch (near heavily diseased tree) | Healthy appearing leaf | F | - | +++ | ITC-A | nt |
| **144** | Vartiokylä (VTKR2) | Large healthy birch (near heavily diseased tree) | Healthy appearing leaf | F | - | +++ | ITC-A | nt |
| **145** | Vartiokylä (VTKR2) | Large healthy birch (near heavily diseased tree) | Healthy appearing leaf | F | + | +++ | np | nt |
| **146** | Vartiokylä (VTKR2) | Large healthy birch (near heavily diseased tree) | Healthy appearing leaf | F | - | +++ | ITC-I | nt |
| **147** | Vartiokylä | Large birch broom like tumors no shoot elongation from tumors (VTKS1). | Broom tree leaf with chlorotic regions and gray spots | G | - | +++ | ITC-F | nt |
| **148** | Vartiokylä | Large birch broom like tumors no shoot elongation from tumors (VTKS1). | Broom tree leaf with chlorotic regions and gray spots | G | - | +++ | ITC-A | nt |
| **149** | Vartiokylä | Large birch broom like tumors no shoot elongation from tumors (VTKS1). | Broom tree leaf with chlorotic regions and gray spots | G | - | +++ | ITC-A | nt |
| **150** | Vartiokylä | Large birch broom like tumors no shoot elongation from tumors (VTKS1). | Broom tree leaf with chlorotic regions and gray spots | G | - | +++ | ITC-F | nt |
| **151** | Vartiokylä | Large birch broom like tumors no shoot elongation from tumors (VTKS1). | Broom tree leaf with chlorotic regions and gray spots | G | - | +++ | ITC-D | RGR-2 |
| **152** | Vartiokylä | Large birch broom like tumors no shoot elongation from tumors (VTKS1). | Broom tree leaf with chlorotic regions and gray spots | G | ++ | +++ | ITC-F | nt |
| **153** | Vartiokylä | Large birch broom like tumors no shoot elongation from tumors (VTKS1). | Broom tree leaf with chlorotic regions and gray spots | G | - | +++ | ITC-F | nt |
| **154** | Vartiokylä | Large birch broom like tumors no shoot elongation from tumors (VTKS1). | Broom tree leaf with chlorotic regions and gray spots | G | - | +++ | ITC-F | nt |

| | | | | | | | | |
|---|---|---|---|---|---|---|---|---|
| **155** | Vartiokylä | Large birch broom like tumors no shoot elongation from tumors (VTKS1). | Broom tree leaf with chlorotic regions and gray spots | G | - | +++ | ITC-F | nt |
| **156** | Vartiokylä | Large birch broom like tumors no shoot elongation from tumors (VTKS1). | Broom tree leaf with chlorotic regions and gray spots | G | - | +++ | ITC-F | nt |
| **157** | Vartiokylä | Large birch broom like tumors no shoot elongation from tumors (VTKS1). | Broom tree leaf with chlorotic regions and gray spots | G | - | +++ | ITC-F | nt |
| **158** | Vartiokylä | Large birch broom like tumors no shoot elongation from tumors (VTKS1). | Broom tree leaf with chlorotic regions and gray spots | G | - | +++ | ITC-F | nt |
| **159** | Vartiokylä | Large birch broom like tumors no shoot elongation from tumors (VTKS1). | Broom tree leaf with chlorotic regions and gray spots | G | + | + | ITC-E | nt |
| **160** | Vartiokylä | Large birch broom like tumors no shoot elongation from tumors (VTKS1). | Broom tree leaf with chlorotic regions and gray spots | G | - | +++ | ITC-A | nt |
| **161** | Vartiokylä | Large birch broom like tumors no shoot elongation from tumors (VTKS1). | Broom tree leaf with chlorotic regions and gray spots | G | +++ | +++ | np | nt |
| **162** | Vartiokylä | Large birch broom like tumors no shoot elongation from tumors (VTKS1). | Broom tree leaf with chlorotic regions and gray spots | G | - | +++ | ITC-A | nt |
| **163** | Vartiokylä | Large birch broom like tumors no shoot elongation from tumors (VTKS1). | Broom tree leaf with chlorotic regions and gray spots | G | - | +++ | ITC-E | nt |
| **164** | Vartiokylä | Large birch broom like tumors no shoot elongation from tumors (VTKS1). | Broom tree leaf with chlorotic regions and gray spots | G | + | +++ | ITC-E | nt |
| **165** | Vartiokylä | Large birch broom like tumors no shoot elongation from tumors (VTKS1). | Broom tree leaf with chlorotic regions and gray spots | G | - | +++ | ITC-E | nt |
| **166** | Vartiokylä | Large birch broom like tumors no shoot elongation from tumors (VTKS1). | Broom tree leaf with chlorotic regions and gray spots | G | - | - | np | nt |
| **167** | Vartiokylä | Large birch broom like tumors no shoot elongation from tumors (VTKS1). | Broom tree leaf with chlorotic regions and gray spots | G | +++ | +++ | np | nt |
| **168** | Herttoniemi | HERS1 Large planted birch lots of elongated small brooms | Broom tree leaf with gray spots | H | - | +++ | ITC-A | nt |
| **169** | Herttoniemi | HERS1 Large planted birch lots of elongated small brooms | Broom tree leaf with gray spots | H | - | +++ | ITC-A | nt |
| **170** | Herttoniemi | HERS1 Large planted birch lots of elongated small brooms | Broom tree leaf with gray spots | H | - | +++ | ITC-A | nt |
| **171** | Herttoniemi | HERS1 Large planted birch lots of elongated small brooms | Broom tree leaf with gray spots | H | - | ++ | ITC-E | nt |
| **172** | Herttoniemi | HERS1 Large planted birch lots of elongated small brooms near to gate | Broom tree leaf with gray spots | H | - | ++ | ITC-F | nt |
| **173** | Herttoniemi | HERS1 Large planted birch lots of elongated small brooms | Broom tree leaf with gray spots | H | - | +++ | ITC-A | nt |
| **174** | Herttoniemi | HERS1 Large planted birch lots of elongated small brooms | Broom tree leaf with gray spots | H | + | +++ | ITC-A | nt |
| **175** | Herttoniemi | HERS1 Large planted birch lots of elongated small brooms | Broom tree leaf with gray spots | H | - | +++ | ITC-A | nt |
| **176** | Herttoniemi | HERS1 Large planted birch lots of elongated small brooms | Broom tree leaf with gray spots | H | - | +++ | ITC-A | nt |
| **177** | Herttoniemi | HERS1 Large planted birch lots of elongated small brooms | Broom tree leaf with gray spots | H | - | +++ | ITC-A | nt |
| **178** | Herttoniemi | HERS1 Large planted birch lots of elongated small brooms | Broom tree leaf with gray spots | H | + | +++ | ITC-A | nt |
| **179** | Herttoniemi | HERS1 Large planted birch, many elongated small brooms | Broom tree leaf with gray spots | H | - | +++ | ITC-A | nt |

| | | | | | | | | |
|---|---|---|---|---|---|---|---|---|
| **180** | Herttoniemi | HERS1 Large planted birch many elongated small brooms | Broom tree leaf with gray spots | H | - | +++ | ITC-F | nt |
| **181** | Herttoniemi | HERS1 Large planted birch lots of elongated small brooms | Broom tree leaf with gray spots | H | - | +++ | ITC-A | nt |
| **182** | Herttoniemi | HERS1 Large planted birch lots of elongated small brooms | Broom tree leaf with gray spots | H | - | +++ | ITC-A | nt |
| **183** | Herttoniemi | HERS1 Large planted birch lots of elongated small brooms | Broom tree leaf with gray spots | H | - | +++ | ITC-F | nt |
| **184** | Herttoniemi | HERS1 Large planted birch lots of elongated small brooms | Broom tree leaf with gray spots | H | - | +++ | ITC-F | nt |
| **185** | Herttoniemi | HERS1 Large planted birch lots of elongated small brooms | Broom tree leaf with gray spots | H | - | ++ | ITC-F | nt |
| **186** | Herttoniemi | HERS1 Large planted birch lots of elongated small brooms | Broom tree leaf with gray spots | H | + | +++ | ui | nt |
| **187** | Herttoniemi | HERS1 Large planted birch lots of elongated small brooms | Broom tree leaf with gray spots | H | - | - | np | nt |
| **188** | Herttoniemi | HERS1 Large planted birch lots of elongated small brooms | Broom tree leaf with gray spots | H | - | ++++ | ui | nt |
| **189** | Herttoniemi | HERS1 Large planted birch lots of elongated small brooms | Broom tree leaf with gray spots | H | - | ++++ | ui | nt |
| **190** | Herttoniemi | HERS2 Large planted birch less elongated small brooms | Broom tree leaf with chlorotic regions and gray spots | I | - | ++++ | ITC-A | nt |
| **191** | Herttoniemi | HERS2 Large planted birch less elongated small brooms | Broom tree leaf with chlorotic regions and gray spots | I | - | ++++ | ITC-A | nt |
| **192** | Herttoniemi | HERS2 Large planted birch less elongated small brooms | Broom tree leaf with chlorotic regions and gray spots | I | - | ++++ | ITC-A | nt |
| **193** | Herttoniemi | HERS2 Large planted birch less elongated small brooms | Broom tree leaf with chlorotic regions and gray spots | I | - | ++++ | ITC-A | nt |
| **194** | Herttoniemi | HERS2 Large planted birch less elongated small brooms | Broom tree leaf with chlorotic regions and gray spots | I | - | +++ | ITC-A | nt |
| **195** | Herttoniemi | HERS2 Large planted birch less elongated small brooms | Broom tree leaf with chlorotic regions and gray spots | I | - | +++ | ITC-A | nt |
| **196** | Herttoniemi | HERS2 Large planted birch less elongated small brooms | Broom tree leaf with chlorotic regions and gray spots | I | - | +++ | ITC-A | nt |
| **197** | Herttoniemi | HERS2 Large planted birch, less elongated small brooms | Broom tree leaf with chlorotic regions and gray spots | I | - | +++ | ITC-C | RGR-0 |
| **198** | Herttoniemi | HERS2 Large planted birch, less elongated small brooms | Broom tree leaf with chlorotic regions and gray spots | I | - | ++ | ITC-D | RGR-3 |
| **199** | Herttoniemi | HERS2 Large planted birch, less elongated small brooms | Broom tree leaf with chlorotic regions and gray spots | I | - | ++ | ITC-D | RGR-2 |
| **200** | Herttoniemi | HERS2 Large planted birch, less elongated small brooms | Broom tree leaf with chlorotic regions and gray spots | I | - | +++ | ITC-S | nt |
| **201** | Herttoniemi | HERS2 Large planted birch, less elongated small brooms | Broom tree leaf with chlorotic regions and gray spots | I | - | +++ | ITC-T | nt |
| **202** | Herttoniemi | HERS2 Large planted birch, less elongated small brooms | Broom tree leaf with chlorotic regions and gray spots | I | - | +++ | ITC-F | nt |
| **203** | Herttoniemi | HERS2 Large planted birch, less elongated small brooms | Broom tree leaf with chlorotic regions and gray spots | I | - | +++ | ITC-A | nt |
| **204** | Herttoniemi | HERS2 Large planted birch, less elongated small brooms | Broom tree leaf with chlorotic regions and gray spots | I | - | +++ | ITC-A | nt |

| | | | | | | | | |
|---|---|---|---|---|---|---|---|---|
| **205** | Herttoniemi | HERS2 Large planted birch, less elongated small brooms | Broom tree leaf with chlorotic regions and gray spots | I | - | +++ | ITC-C | RGR-0 |
| **206** | Herttoniemi | HERS2 Large planted birch, less elongated small brooms | Broom tree leaf with chlorotic regions and gray spots | I | - | +++ | ITC-C | RGR-0 |
| **207** | Herttoniemi | HERS2 Large planted birch, less elongated small brooms | Broom tree leaf with chlorotic regions and gray spots | I | - | ++ | ITC-C | RGR-0 |
| **208** | Herttoniemi | HERS2 Large planted birch, less elongated small brooms | Broom tree leaf with chlorotic regions and gray spots | I | - | ++ | ui | nt |
| **209** | Herttoniemi | HERS2 Large planted birch, less elongated small brooms | Broom tree leaf with chlorotic regions and gray spots | I | - | +++ | ITC-C | RGR-0 |
| **210** | Herttoniemi | HERS2 Large planted birch, less elongated small brooms | Broom tree leaf with chlorotic regions and gray spots | I | - | +++ | ITC-A | nt |
| **211** | Herttoniemi | HERS2 Large planted birch, less elongated small brooms | Broom tree leaf with chlorotic regions and gray spots | I | - | +++ | ITC-D | RGR-2 |
| **212** | Herttoniemi | HERS2 Large planted birch, less elongated small brooms | Broom tree leaf with chlorotic regions and gray spots | I | - | +++ | ITC-C | RGR-0 |
| **213** | Herttoniemi | HERS2 Large planted birch, less elongated small brooms | Broom tree leaf with chlorotic regions and gray spots | I | +++ | +++ | ITC-C | RGR-0 |
| **214** | | HERS2 Large planted birch, less elongated small brooms | Broom tree leaf with chlorotic regions and gray spots | I | - | +++ | ITC-C | RGR-0 |
| **215** | Herttoniemi | HERS2 Large planted birch, less elongated small brooms | Broom tree leaf with chlorotic regions and gray spots | I | - | +++ | ITC-C | RGR-0 |
| **216** | Herttoniemi | HERS2 Large planted birch, less elongated small brooms | Broom tree leaf with chlorotic regions and gray spots | I | +++ | +++ | np | nt |
| **217** | Herttoniemi | HERS2 Large planted birch, less elongated small brooms | Broom tree leaf with chlorotic regions and gray spots | I | - | +++ | ITC-C | RGR-0 |
| **218** | Vartioharju (VTH) | Large birch(hanging phenotype) with elongated brooms, heavily diseased(VTHS1) | Broom leaf gray spots and yellow spot and deficient growth | E | - | +++ | ITC-D | RGR-0 |
| **219** | Vartioharju (VTH) | Large birch(hanging phenotype) with elongated brooms, heavily diseased(VTHS1) | Broom leaf gray spots and yellow spot and deficient growth | E | - | +++ | ITC-D | RGR-2 |
| **220** | Vartioharju (VTH) | Large birch(hanging phenotype) with elongated brooms, heavily diseased(VTHS1) | Broom leaf gray spots and yellow spot and deficient growth | E | - | +++ | ITC-D | RGR-2 |
| **221** | Vartioharju (VTH) | Large birch(hanging phenotype) with elongated brooms, heavily diseased(VTHS1) | Broom leaf gray spots and yellow spot and deficient growth | E | - | ++ | ITC-F | nt |
| **222** | Vartiokylä | Large birch broom like tumors no shoot elongation from tumors (VTKS1). | | G | - | +++ | ui | nt |
| **223** | Herttoniemi | HERS1 Large planted birch lots of elongated small brooms near to gate | Surface sterilized leaf that was chopped | H’ | +++ | +++ | ui | nt |
| **224** | Herttoniemi | HERS1 Large planted birch lots of elongated small brooms near to gate | Surface sterilized leaf that was chopped | H’ | +++ | +++ | ui | nt |

Supplementary table 2. Taphrina betulina strain diversity from different broom morphology types.

| Species | NO BROOM | EB | TL | SE | Total |
|---|---|---|---|---|---|
| *Taphrina betulina* strains | | | | | |
| ITC-C RsaI-0 | | 2 | | 1 | 3 |
| ITC-C RsaI-1 | 1 | | | | 1 |
| ITC-C RsaI-2 | | 1 | | | 1 |
| ITC-D RsaI-0 | 1 | 9 | | | 10 |
| ITC-D RsaI-1 | | 1 | | | 1 |
| ITC-D RsaI-2 | 4 | 23 | 1 | 1 | 29 |
| ITC-D RsaI-3 | | 1 | | 2 | 3 |
| ITC-U | | | | 9 | 9 |
| *Taphrina total* | 6 | 37 | 1 | 13 | **57** |

Supplementary table 3. Cell Size of 22 T. betulina selected strains

| Type | Strains | Cell Size | | | |
|---|---|---|---|---|---|
| | | Length (µm) | Significance group | Width (µm) | Significance group |
| I* | 25 | 4.96±1.1 | ab | 3.39±0.8 | abcde |
| | 26 | 5.07±0.9 | ab | 3.5±0.7 | ab |
| | 31 | 4.75±1.3 | ab | 3.17±0.8 | df |
| | 34 | 4.9±0.9 | ab | 3.24 ±0.7 | abce |
| | 85 | 4.82±0.9 | ab | 3.51±0.7 | abcdef |
| | 112 | 5.12±0.9 | ab | 3.61±0.8 | abce |
| | 219 | 5.13 ±0.8 | ab | 3.5 ±0.7 | abcdef |
| II** | 58 | 5.79±1.2 | ac | 3.58±0.8 | acdef |
| | 59 | 5.08±1.1 | ab | 3.7±0.9 | abcdef |
| | 62 | 4.95±1 | ab | 3.5±0.7 | abcde |
| | 63 | 4.81±0.9 | ab | 3.33±0.6 | cdef |
| | 68 | 4.96±0.87 | b | 3.45±0.7 | b |
| | 69 | 5.02±1.1 | ab | 3.39±0.6 | abc |
| | 82 | 5±1 | d | 3.41±0.7 | cdef |
| | 83 | 4.87±1 | ab | 3.27±0.7 | df |
| | 151 | 5.16±1.1 | ab | 3.4±0.7 | abcde |
| | 198 | 5.15±0.9 | ab | 3.69±0.8 | abcde |
| | 199 | 4.81±0.9 | ab | 3.3±0.6 | abde |
| III*** | 11 | 5.65±1.2 | cd | 3.81±1.2 | f |
| | 19 | 4.88±1 | ab | 3.22±0.7 | def |
| | 20 | 5.28±1 | ab | 3.56±0.7 | b |
| | 129 | 5.08±1.1 | ab | 3.18±0.6 | abcdef |
